# The Brain Electroencephalogram Microdisplay for Precision Neurosurgery

**DOI:** 10.1101/2023.07.19.549735

**Authors:** Youngbin Tchoe, Tianhai Wu, Hoi Sang U, David M. Roth, Dongwoo Kim, Jihwan Lee, Daniel R. Cleary, Patricia Pizarro, Karen J. Tonsfeldt, Keundong Lee, Po Chun Chen, Andrew M. Bourhis, Ian Galton, Brian Coughlin, Jimmy C. Yang, Angelique C. Paulk, Eric Halgren, Sydney S. Cash, Shadi A. Dayeh

## Abstract

Brain surgeries are among the most delicate clinical procedures and must be performed with the most technologically robust and advanced tools. When such surgical procedures are performed in functionally critical regions of the brain, functional mapping is applied as a standard practice that involves direct coordinated interactions between the neurosurgeon and the clinical neurology electrophysiology team. However, information flow during these interactions is commonly verbal as well as time consuming which in turn increases the duration and cost of the surgery, possibly compromising the patient outcomes. Additionally, the grids that measure brain activity and identify the boundaries of pathological versus functional brain regions suffer from low resolution (3-10 mm contact to contact spacing) with limited conformity to the brain surface. Here, we introduce a brain intracranial electroencephalogram microdisplay (Brain-iEEG-microdisplay) which conforms to the brain to measure the brain activity and display changes in near real-time (40 Hz refresh rate) on the surface of the brain in the surgical field. We used scalable engineered gallium nitride (GaN) substrates with 6” diameter to fabricate, encapsulate, and release free-standing arrays of up to 2048 GaN light emitting diodes (μLEDs) in polyimide substrates. We then laminated the μLED arrays on the back of micro-electrocorticography (μECoG) platinum nanorod grids (PtNRGrids) and developed hardware and software to perform near real-time intracranial EEG analysis and activation of light patterns that correspond to specific cortical activities. Using the Brain-iEEG-microdisplay, we precisely ideFSntified and displayed important cortical landmarks and pharmacologically induced pathological activities. In the rat model, we identified and displayed individual cortical columns corresponding to individual whiskers and the near real-time evolution of epileptic discharges. In the pig animal model, we demonstrated near real-time mapping and display of cortical functional boundaries using somatosensory evoked potentials (SSEP) and display of responses to direct electrical stimulation (DES) from the surface or within the brain tissue. Using a dual-color Brain-iEEG-microdisplay, we demonstrated co-registration of the functional cortical boundaries with one color and displayed the evolution of electrical potentials associated with epileptiform activity with another color. The Brain-iEEG-microdisplay holds the promise of increasing the efficiency of diagnosis and possibly surgical treatment, thereby reducing the cost and improving patient outcomes which would mark a major advancement in neurosurgery. These advances can also be translated to broader applications in neuro-oncology and neurophysiology.

**One Sentence Summary:** A brain intracranial electroencephalogram microdisplay (Brain-iEEG-microdisplay) measures and displays real-time brain activity in the surgical field.

## INTRODUCTION

One of the most important goals of brain tumor treatment is complete surgical resection of the pathologic tissue so as to improve survival rate. However, in neuro-oncology and surgical interventions for epilepsy, there is usually a trade-off between the extent of resection and the risk of postoperative neurological deficit due to loss of functional tissue.(*1–6*) Furthermore, a brain lesion can distort normal anatomy and cause functional reorganization(*7*) that may not be fully identified with non-invasive functional neuroimaging techniques such as positron emission tomography (PET), functional magnetic resonance imaging (fMRI), and Magneto-encephalography (MEG). Therefore, direct electrical stimulation (DES) paired with electrophysiological recording has evolved as the gold standard for determining functional boundaries before resection of lesions located in eloquent areas.(*8–12*) DES is a reliable, safe, and effective technique for the near real-time precise identification and preservation of cortical and subcortical neuronal pathways for motor and sensory function, language, and even memory.(*8, 12*) More recently, passive electrophysiological recordings in the high gamma band (60-200 Hz) are shown to be a fast, robust, and reliable method for identifying receptive language areas(*13*) with differences in brain activity entropy and high gamma activity characterizing pathological tissue in brain tumors.(*3, 14, 15*) Based on DES mapping, functional regions are marked and preserved from resection, usually with a margin of about 5 mm around the motor areas and 7 mm around the language areas to avoid post-operative functional deficit.(*16*) Most commonly, identifying the spatial extent of functional and pathological boundaries based on neurophysiology is communicated verbally by messages written on paper that is transported across the operating room between the intraoperative monitoring team comprised of clinical electrophysiologists and neurologists and the neurosurgeons. Moreover, some groups use sterile paper placed on the surface of the brain to mark areas for resection and areas for preservation.(*17, 18*) Clearly, the resolution and the methodology for presenting this critical information to guide neurosurgical interventions can be significantly improved.

Recent advances in thin-film microelectrode technology have increased the resolution by which cortical activity can be measured and localized.(*19–21*) For example, the platinum nanorod grids (PtNRGrids) have been used to identify the curvilinear boundary between functional and pathological tissues in the human brain at unprecedented spatial resolution.(*21*) However, to date, there is no available technology that can display these cortical boundaries in situ on the brain. The rise of flexible panel display technology presents an opportunity to revolutionize the way we measure and display cortical activity. In the display technology field, gallium nitride (GaN) light emitting diodes (LEDs) represent an efficient and scalable solution for solid-state lighting and displays,(*22*) and have recently progressed to full-color capability with layer-transfer techniques of micro-LEDs (μLEDs).(*23*) However, the monolithic integration, encapsulation, and release of free-standing GaN μLED arrays from wafers suitable for production scale with diameters exceeding 6” has not been previously accomplished.

To address the need for high fidelity, near real-time tracking of functional and pathological brain activity as well as advance the use of powerful new GaN μLED display tools, here we describe a novel fabrication procedure for flexible GaN μLEDs and the application of this innovation to a flexible brain electroencephalogram micro-display (Brain-iEEG-microdisplay). The Brain-iEEG-microdisplay comprises GaN μLEDs mounted behind the PtNRGrids, as well as acquisition and control electronics and software drivers to analyze and project the cortical activity directly from the surface of the brain. We demonstrated the safe and effective use of the Brain-iEEG-microdisplay through benchtop characterization. Moreover, we demonstrated highly localized brain activity could be projected via the light display in the Brain-iEEG-microdisplay in animal experiments in rats and pigs. In the pig model, we demonstrated the capabilities of the Brain-iEEG-microdisplay to provide automated and near real-time visual representation of somatosensory evoked potentials (SSEPs). In addition, large voltage deflections correlated with epileptic discharges were co-registered to mapped sensory regions using dual-color Brain-iEEG-microdisplay. In the rat model, we showed that the Brain-iEEG-microdisplay can resolve and represent individual cortical columns and can display pathological activity with high definition. Overall, our Brain-iEEG-microdisplay addresses technological shortcomings during neurosurgical procedures and can provide fine resection boundaries from the brain surface at high spatial and temporal resolution (Fig. 1A).

**Figure 1.**
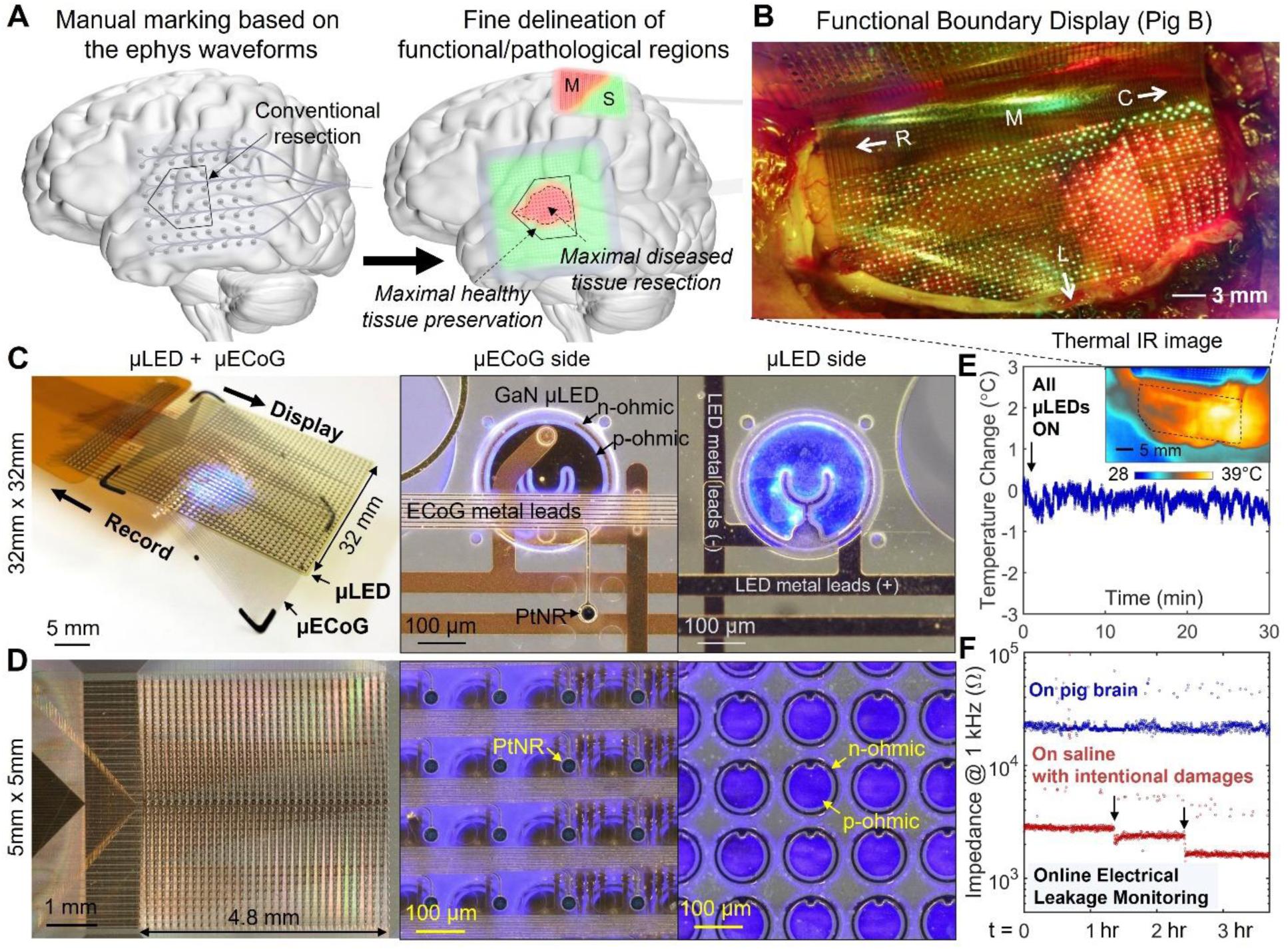
Innovation in the way brain surgery is conducted by a Brain-iEEG-microdisplay to inform high-precision brain surgery. (A) Core concept of the Brain-iEEG-microdisplay that simultaneously measures and displays in near real-time healthy and/or diseased brain regions and boundaries directly from the surgical field. (B) Delineation of the curvilinear boundary between the motor and sensory cortical regions on the pig’s brain using a dual-color Brain-iEEG-display (image taken under a 550 nm blue light blocking lens). Device structures and magnified OM images of (C) 32mm x 32mm and (D) 5mm x 5mm μLED+μECoG devices that integrated the flexible μLED arrays with 1024 pixels and PtNRGrids with 1024 channels. (E) Thermal safety evaluation of the Brain-iEEG-microdisplay by monitoring the cortical surface temperature of the pig brain under 30 min continuous operation of all the LEDs. (F) Electrical safety evaluation of Brain-iEEG-microdisplay by monitoring the impedance of the device with respect to the brain tissues over 3.7 hours on the pig’s brain.

## RESULTS

### Integration of the Brain-iEEG-microdisplay

To construct the Brain-iEEG-microdisplay, we combined our ultrathin microelectrocorticography (μECoG) grids, the PtNRGrids, with GaN μLEDs. We chose GaN μLEDs because they are highly efficient and consume low power(*23, 24*) thereby producing low thermal heating losses and inducing minimal heating effects on the cortical tissue.(*25*) Additionally, the high intensity of light produced by GaN μLEDs is visible to the human eye even under the bright surgical lightning.(*26*) To fabricate flexible GaN μLED arrays for both high resolution and broad spatial coverage, we developed a scalable process on 6-inch GaN/polycrystalline AlN engineered engineered Qromis® Substrate Technology™ (QST) substrate(*27, 28*) and released an ultrathin μLED arrays, some with 1024 and some with 2048 pixels, embedded in polyimide layers (see Fig. S1). GaN μLEDs with indium gallium nitride (InGaN) quantum wells emitted blue light with peak wavelength of 450 nm (Fig. S2). In order to simultaneously display and co-register normal and diseased cortical activity, we used ink-jet printing to deposit quantum dot color conversion (QDCC) inks made of indium phosphide (InP) quantum dots on the surface of the GaN μLEDs (Fig. S2).(*29*) This allowed us to emit multiple colors, enabling richer and more nuanced visual representation of neural activity patterns (Figs. 1B). The flexible GaN μLED arrays were then laminated on the back of the PtNRGrids grids (Fig. 1C)(*21*) to form the EEG-microdisplay. The PtNRGrids were constructed on thin and brain-conformal parylene C substrates with thousands of channels which reliably recorded the spatiotemporal dynamics of brain activity at high-resolution and with broad coverage.(*21, 30*) As both the flexible μLED array and μECoG were based on scalable, and reconfigurable manufacturing processes, we matched their pitch and coverage for a single Brain-iEEG-microdisplay. Here, we discuss results based on Brain-micro-EEG display with 32mm × 32mm coverage and 1mm pitch suitable for human-scale Brain-iEEG-microdisplay and used on the pig brain (Fig. 1C), and 5mm × 5mm coverage with 0.15mm pitch for rat brains (Fig. 1D). The single-color Brain-iEEG-microdisplay was composed of 1024 PtNR recording contacts and 1024 GaN μLED pixels. The dual color Brain-iEEG-microdisplay comprised 2048 QDCC-printed GaN μLEDs with 0.4mm vertical and 0.5mm horizontal pitches, and a corresponding PtNRGrid with 1024 contacts with 0.8mm vertical and 0.5mm horizontal pitches, both providing a 12.8mm × 32mm brain coverage (Fig. 1B). The μECoG grids utilized 30 μm diameter PtNR contacts with an average impedance of 30kΩ at 1kHz. The size of GaN μLED die varied depending on their intended use in Brain-iEEG-microdisplays. Specifically, GaN μLEDs with a diameter of 220μm were optimized for 1 mm pitch Brain-iEEG-microdisplay (Fig. 1C), while those with a diameter of 100μm were designed for Brain-iEEG-microdisplays with higher densities such as that used for the rat (Fig. 1D) or dual-color 2048 μLEDs used for the pig (Fig. 1B).

Light emission due to injection of electrical current in LEDs has a limited efficiency where part of the electrical power supplied to them is lost as thermal energy. This can increase the cortical surface temperature change which must be kept below 1°C.(*31, 32*) Therefore, robust and reliable measures to limit the temperature rise and to ensure detection of any leakage currents in case of failure of the insulation materials for the GaN μLEDs must be taken. To this end, we have characterized the thermal and electrical safety of the Brain-iEEG-microdisplay. In one experiment, we turned on all 2048 μLEDs at their maximum brightness possible with our LED driver chip while the Brain-iEEG-microdisplay was placed on top of the pig’s brain for 30 minutes. The temperature was then monitored with an infrared (IR) imaging camera. No temperature changes within the measurement resolution of the IR camera (0.1 °C) were observed (Fig. 1E). Additionally, post-mortem histochemical analysis on the same pig’s brain showed undetectable differences in the structure of the cortical surface under the Brain-iEEG-microdisplay and the contralateral region (Fig. S3, N=1): The top cortical layer was of comparable thickness, the neuronal density was similar, and the neuron shape was normal. However, when the same experiment was conducted on a rat’s brain with the high-density Brain-iEEG-microdisplay whose pixel pitch was 0.15mm, we observed a temperature increase of up to 6°C in less than 5 minutes when all 1,024 μLEDs were turned on at maximum brightness. We therefore conducted a series of experiments to determine if this temperature change could be minimized by adjusting the duty cycle of the μLEDs. We achieved acceptable brightness through these adjustments with less than 1°C increase in temperature (Fig. S4).

Next, to continuously evaluate the integrity of the insulation materials for the μLEDs and impose preventive measures to shut down the Brain-iEEG-microdisplay upon insulation material degradation or failure, the impedance of all the electrically conducting elements (metal traces, Ohmic contacts, and the GaN) of the μLEDs in relation to the brain tissue was monitored every 10 seconds throughout the entire 3.7-hour duration of pig brain recording. This test revealed no detectable changes in the impedance, indicating that no electrical leakage paths developed for the duration of the experiment (Fig. 1F and Fig. S5). In order to test the validity of our impedance monitoring setup and to ensure it properly represents electrical leakage paths, scratches were purposedly made on the surface of the μLED array. In this setting, a clear drop in impedance was observed whenever there was mechanical or electrical damage to the device. This result indicates that our impedance monitoring setup was reliable in detecting the presence of electrical leakage paths in the Brain-iEEG-microdisplay though continuous real-time monitoring of the actual current changes would be most appropriate for this application.

### The Brain-iEEG-microdisplay projects the M1/S1 Boundary from the Surface of the Brain

The precise localization of the central sulcus, the anatomical boundary between primary motor (M1) and somatosensory (S1) cortices, is crucial in tumor and epileptogenic tissue resections in these brain regions. This boundary is identified at the point where somatosensory evoked potentials (SSEPs) in response to peripheral nerve stimulation exhibit opposite polarity in their potentials commonly recognized by either a negative phase (P20) and the positive phase (N20).(*33, 34*) Commonly, the presence of pathological tissue can shift this functional boundary from its anatomical organization.(*7, 35*) The Brain-iEEG-microdisplay can directly project the M1/S1 boundary from the surface of the brain based on physiological signatures. We applied the 32mm × 32mm Brain-iEEG-microdisplay across the mid-line of the pigs’ brain (N=5 pigs) covering both the left and right primary motor cortex (M1) and primary somatosensory cortex (S1). To evoke SSEPs, we stimulated the peripheral nerves with subdermal twisted pair needles placed in the pig’s left and right forelimbs (Fig 2A, 10mA, 1ms biphasic pules). A small stimulus artifact prior to high amplitude SSEPs were observed within 10∼50ms post-stimulation and exhibited phase reversal indicated by the red arrow-heads of Fig. 2B.(*36*) Our Brain-iEEG-microdisplay revealed a cortical functional boundary (FB) between the primary somatomotor (M1) cortex and primary somatosensory (S1) cortex depicted with a line of illuminated μLEDs (Figs 2C,F) or represented by a change of color with the dual-color areal illuminated μLEDs (Figs. 2I,J). When the Brain-iEEG-microdisplay covered both hemispheres of the pig’s brain (Fig. 2C-H), we observed contralateral evoked SSEPs and correspondent depiction of the FB with respect to the stimulated forelimb. The FB displayed in near real-time with the Brain-iEEG-microdisplay was also in excellent agreement with the offline analysis done with the P20-N20 potential mapping (Figs. 2D, G, Fig. S6) and correlation coefficient mapping (Fig. 2E,H).

**Figure 2.**
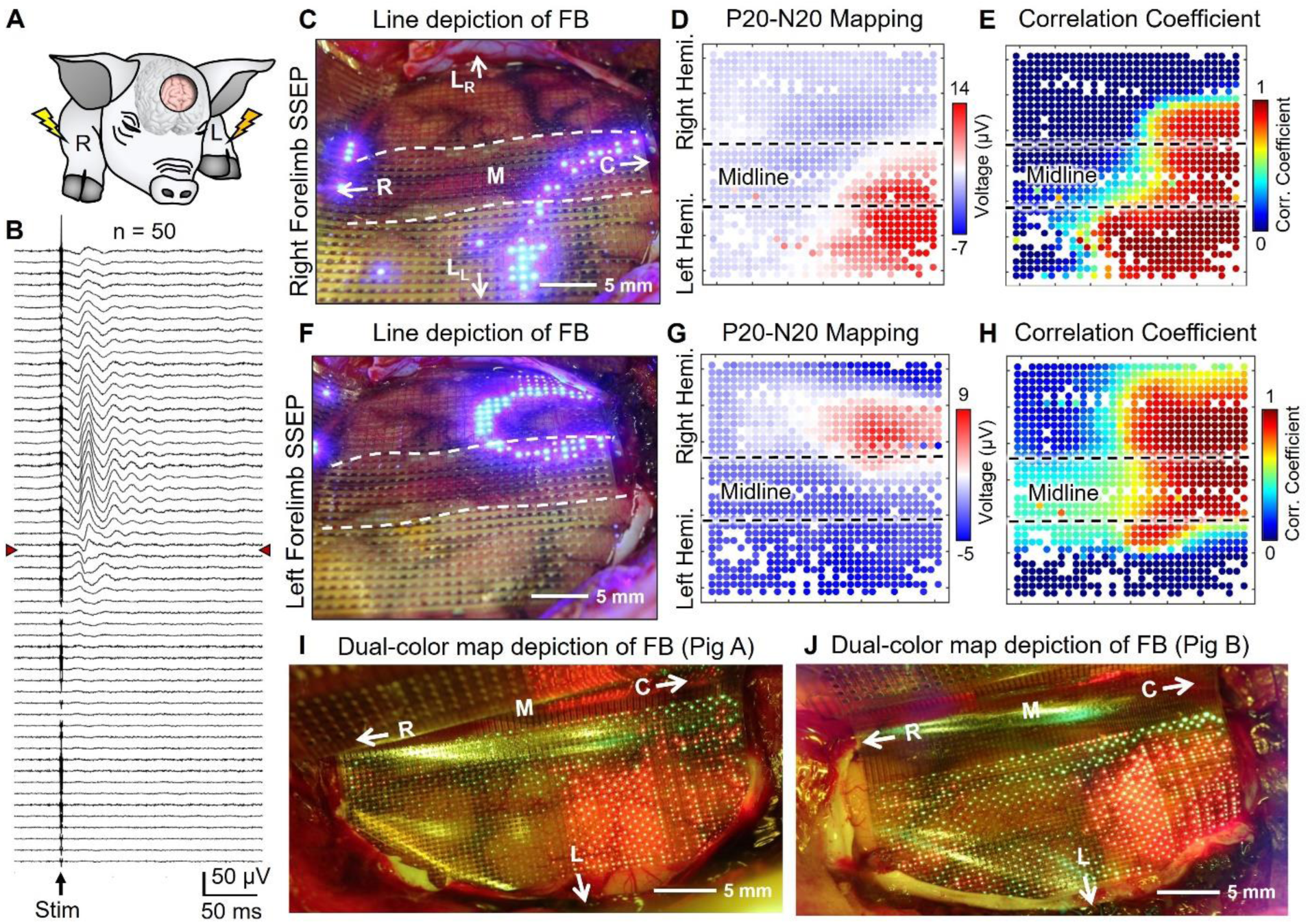
Near real-time mapping and display of the primary somatomotor cortex (M1) and primary somatosensory cortex (S1) tissue landmark for brain surgery in the pig’s brain. (A) Schematics of the experiment depicting the electrical stimulation of the pig’s left and right forelimbs (biphasic, bipolar pulse: 10mA, 1ms) while simultaneously recording SSEPs and displaying the M1/S1 boundary on the surface of the brain. (B) SSEP waveforms measured across an individual row of PtNR contacts across the M1/S1 boundary. The red arrow-heads indicate the phase reversal boundary. (C) Line depiction of M1/S1 functional boundary with the Brain-iEEG-microdisplay using right-forelimb stimulation. The corresponding offline analysis results showing (D) P20-N20 potential and (E) correlation coefficient mapping. (F) Line depiction of M1/S1 functional boundary with the Brain-iEEG-microdisplay using left-forelimb stimulation and (G) corresponding offline analysis of P20-N20 potential and (H) correlation coefficient mapping. Dual-color display (image taken under 550 nm blue light blocking lens) of motor/sensory boundary observed from two different animals: (I) Pig A and (J) Pig B. (C)-(H) Single color (blue) 1024ch Brain-iEEG-microdisplay for M1/S1 boundary. (I)-(J) Dual color (green/red) 2048 ch Brain-iEEG-microdisplay for M1/S1 regions and boundary.

### Brain-iEEG-microdisplay of Functional Cortical Columns from the Surface of the Brain

To demonstrate the advanced mapping and display capabilities of the Brain-iEEG-microdisplay, we conducted a localized sensory mapping on an anesthetized pig by administering electrical or air-puff sensory stimuli to different areas of the pig’s face and limbs thereby evoking high gamma responses on the brain’s surface (Fig 3A). The high gamma activity (HGA) was mapped for each of n=50 trials, and the trial averaged HGA mapping was updated and displayed – every 1 s – on the brain’s surface whenever a new stimulus was administered. Skin electrical stimulation (2mA, 1ms, biphasic, bipolar) of left and right forelimbs, cheeks, and tongue was administered to reveal uniquely distinguishable cortical regions while being able to observe the anatomical features of the brain tissue underlying the Brain-iEEG-microdisplay (Fig 3B-G). We hypothesized that local air-puff stimulation on the surface of the skin delivered through a microcapillary tube can lead to focal responses when compared to gross-sum activation of nerve fibers via electrical stimulation in the vicinity of the stimulating needle probes.(*37*) Indeed, with air-puff stimulation of the tongue tip, much more tightly localized HGA responses were observed (Fig. 3H) compared to responses to nerve electrical stimulation (Fig. 3G). These results are corroborated by responses of electrical and air-puff stimulations of the pig’s snout (Fig. 3I) depicted in waveforms (Figs. 3E,3I) and the spatial mapping of the HGA (Figs. 3J-L). The potential evoked by air-puff stimulation of two different snout positions (Fig. 3I) showed a single peak near 35ms whereas the electrically stimulated snout showed multiple peaks from 10 to 90ms, the latter presumably due to the volumetric stimulation of multiple different types of sensory and motor neurons with varying latencies.(*38*) The Brain-iEEG-microdisplay depicted relatively broader spatial maps of HGA responses evoked by electrical stimulation of the snout (Fig. 3J) and tightly localized HGA by air-puff stimulation of two different spots (Figs. 3K,L) within the electrically responsive region of Fig. 3J. These HGA mappings displayed on the brain surface were in remarkable agreement with the offline analysis (Figs. 3M,N) that shows spatial mapping of HGA together with trial-averaged raw waveforms (n=50 trials).

**Figure 3.**
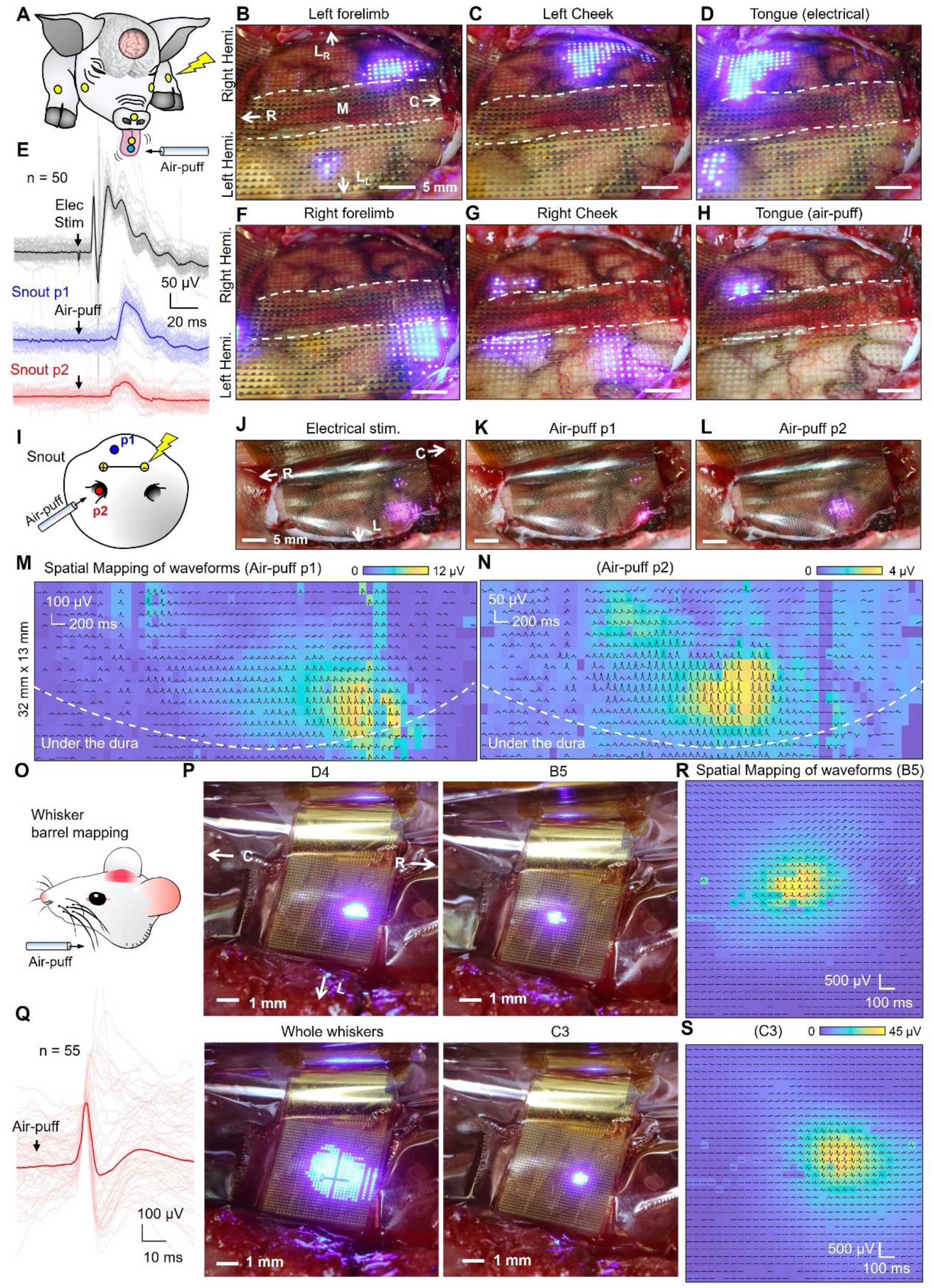
Advanced and fine scale sensory mapping with the Brain-iEEG-microdisplay from the pig and of the rat brains. (A) Schematics showing the locations and type of sensory stimuli on the pig. High gamma activity (HGA) mapping of the electrical stimulation evoked responses (n=50 trials) of (B) left forelimb, (C) left cheek, (D) tongue, (F) right forelimb, and (G) right cheek. (H) HGA mapping evoked by air-puff stimulation of tongue tip. (i) Schematics of electrical and air-puff stimulations of pig snout. (E) Trial average SSEP waveforms (n=50 trials) overlaid with waveforms of individual trials under the electrical and air-puff stimulation of snout. HGA mapping evoked by (J) electrical stimulation of snout and air-puff stimulation of snout at positions (K) p1 and (L) p2. Corresponding spatial mapping of HGA and trial-averaged raw waveforms (n=50 trials) evoked by air-puff of snout at (M) p1 and (N) p2 positions. (O) Schematics of air-puff stimulation of rat whiskers. (P) μLED indicating the individual cortical column position of B5, C3, and D4 barrel cortices by the online trial averaged HGA mapping (n=55 trials). Overall area of barrel cortices was displayed by the multiple whisker stimulation. (Q) Trial average SSEP waveforms (n=55 trials) overlaid with waveforms of individual trials under the air-puff stimulation of whisker. Spatial mapping of HGA and trial averaged waveforms (n=55 trials) by the air-puff stimulation of (R) B5 and (S) C3 whiskers.

To demonstrate fine scale detection and projection of brain activity to the µLEDs display, we applied the 5mm × 5mm high-resolution Brain-iEEG-microdisplay to the barrel cortex of the rat to isolate the location and boundaries of sub-millimeter-scale cortical columns. The rat barrel cortex is a well-studied system where specific sensory cortical columns have a one-to-one mapping with individual whiskers.(*39*) While air-puffs stimulated the individual whiskers, we recorded and displayed the spatial map of trial averaged (n=55 trials, N=4 rats) HGA. We observed that a small group of µLEDs were selectively lit up in clearly distinct positions at sub-millimeter scale indicating individual positions of cortical columns that correspond to each whisker (Fig. 3P). When all whiskers in the field-of-blow were air-puff stimulated, a larger group of µLEDs were lit up with a diameter near 3 mm, indicative of the extent of multiple whisker barrel cortices (Fig. 3P). When compared with the spatial mapping by offline analysis of recorded waveforms (Fig. 3Q), the spatial mapping of raw waveforms and HGA (n=55 trials) (Fig. 3R,S) were in excellent agreement with the HGA displayed with Brain-iEEG-microdisplay.

### The Brain-iEEG-microdisplay Informs the Extent of Clinical Stimulation

Direct electrical stimulation of the brain and observation of behavioral responses and after-discharges inform the location and boundaries of diseased tissue and is typically carried out with a handheld bipolar cortical stimulator (Ojemann probe).(*40*) This procedure is generally performed on the exposed brain without any grid on the surface. Our Brain-iEEG-microdisplay was designed with large-diameter through-hole arrays (0.5 mm diameter with 1 mm pitch) throughout the grid (Fig. S7). These perforations permitted the delivery of stimulation with the Ojemann probe to any position of the brain together with the simultaneous display of the extent of the resultant electrical potential and the brain’s response. We administered charge-balanced biphasic pulses of 3 mA and 50 Hz with varying durations between 0.1 s to 1.9 s and recorded and displayed the root mean square (RMS) of the voltage brain wave responses within the frequency range of 10-59 Hz. The refresh rate of the Brain-iEEG-microdisplay was 40 Hz as we processed 25ms data packets from 1024 channels and displayed the RMS map every 25ms. During brain stimulation, a large electrical potential was created causing many µLEDs in the array to light up around each side of bipolar probe (Fig. 4B). After the stimulation, we observed a post-stimulus pattern around the bipolar probe near 200 ms (Fig. 4C). The RMS potential map observed in near real-time on the brain surface was in good agreement with the offline analysis where the spatial map of raw waveforms was overlaid with the RMS potential map (Figs. 4D and E).

**Figure 4.**
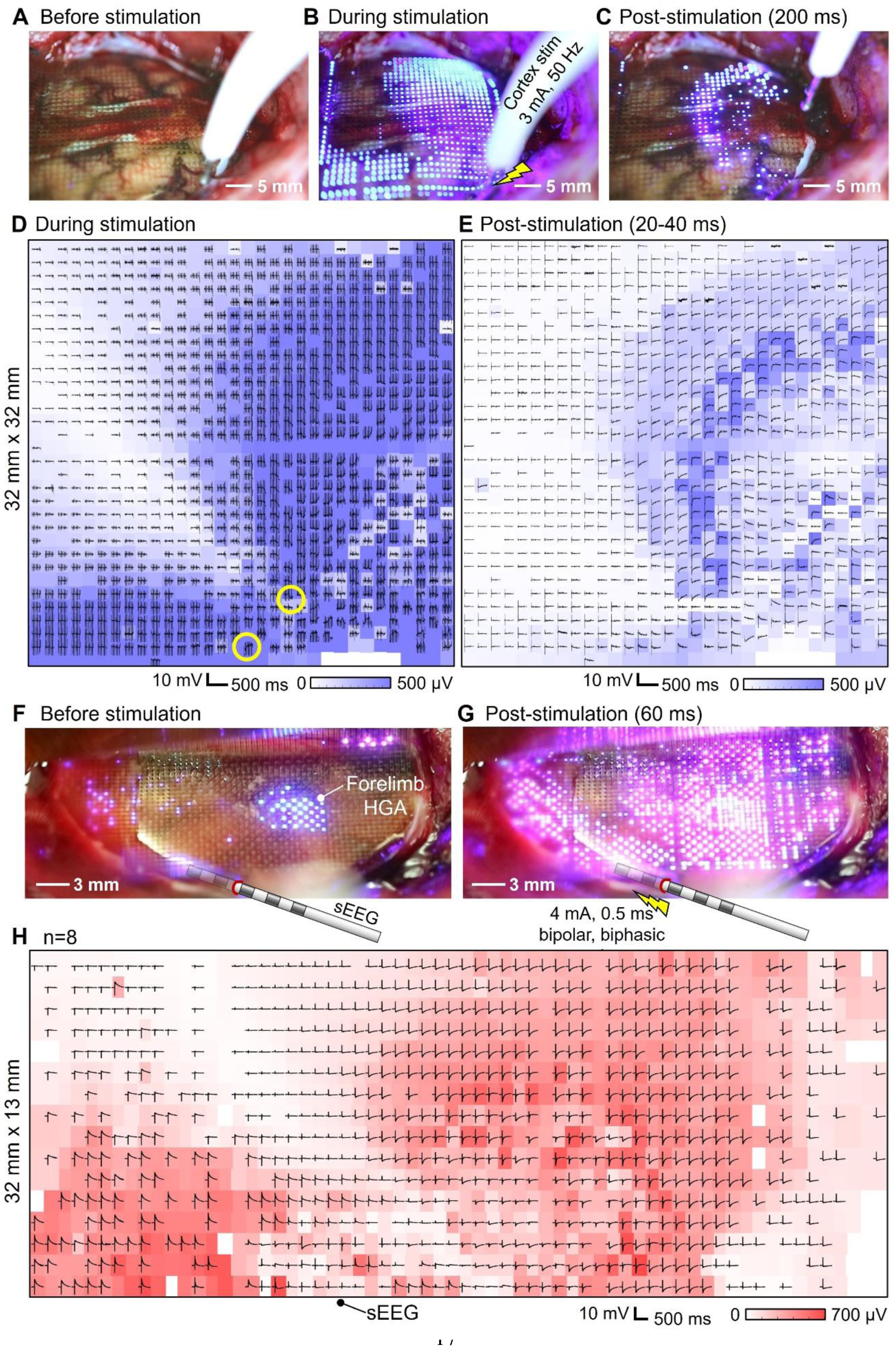
The Brain-iEEG-microdisplay allows visualization of electrical potentials resulting from direct electrical stimulation with clinical probes. Map of the electrical potential created on the brain surface (A) before, (B) during the electrical stimulation (3mA, 50 Hz) and (C) post stimulation (200 ms). Offline analysis of raw waveforms and RMS potential map (10-59 Hz) (D) during the stimulation (-200∼0 ms) and (E) after the stimulation (-10 ∼ 200 ms); the RMS potential map for (E) was obtained between 20-40 ms post-stimulus. Potential map displayed on the cortical surface with (F) ‘blue’ color showing the HGA corresponding to forelimb and (G) ‘pink’ color showing the online potential map evoked by the 4 mA electrical stimulations by sEEG electrode between the pair of electrodes at the depth of 9mm and 3mm. (H) Trial-averaged (n=8 trials) waveform mapping (-100 ∼ 300 ms) together with RMS potential map (35-60 ms, 10-59 Hz) after 4 mA single (left) and train (right) of pulses. (E) Trial averaged waveforms of sEEG electrical stimulation evoked responses from a single channel.

In addition to the brain surface Ojemann stimulation, depth electrode stimulation was also used to map corticocortical structures. To this end, we inserted a stereoelectroencephalography (sEEG) probe (0.8 mm diameter, contact spacing of 6 mm) into the depth of the brain (Fig. 4A) next to the Brain-iEEG-microdisplay (see the position marked in Figs. 4D and 4G). We then applied bipolar, biphasic stimulation pulses (4 mA, 0.5 ms, single pulse) between adjacent contacts at depths of 9 mm and 3 mm, respectively. Under a single pulse stimulation, a population of pink LEDs lit up around the sEEG stimulation site indicating a clear spatial display of post-stimulus potentials on the brain surface (Fig. 4G). Here, a dual-color Brain-iEEG-microdisplay with 2048 µLEDs displayed a statically lit HGA corresponding to the forelimb in blue and the near real-time evolution of the RMS potentials in pink (Fig. 4F). The spatial pattern of the responses observed on the Brain-iEEG-microdisplay was in good agreement with the offline analysis which overlaid the post-stimulus RMS potential map (35-60 ms, 10-59 Hz) together with the spatial map of waveforms including the stimulation pulse (-100 ∼ 300 ms) (Fig. 4H). The Brain-iEEG-microdisplay, therefore, provided previously unattainable near real-time feedback of stimulation responses directly from the surface of the brain (Movies S1 and S2).

### The Brain-iEEG-microdisplay Features Near Real-Time Videos of Pathological Wave Dynamics

A final important feature of the Brain-iEEG-microdisplay is its ability to facilitate high-resolution mapping of pathological brain activity. For example, the precise intraoperative neuromonitoring to detect the onset zones of seizures and patterns of their spread is essential for successful treatment.(*41*) To demonstrate pathological brain activity such as epileptiform activity could be displayed using the Brain-iEEG-microdisplay, we placed the display over the anesthetized pig’s brain and artificially induced epileptic seizures by applying neurotoxins on the brain underlying the display (Fig. 5A). This procedure included the administration of three types of neurotoxins, with each toxin administered to separate animals in individual cases: (1) Bicuculline (BIC) crystals that were topically applied on the brain surface (N=1 pig case), (2) the sub-cortical injection of 4-Aminopyrodine (4-AP) (N=1 pig case) or Penicillin-G (Pen-G) (N=2 pig cases). All three types of neurotoxins induced epileptiform activities while BIC induced the most prominent responses showing large amplitude epileptiform activities (∼0.5mV peak-to-peak) (Fig. 5B). After the initial observation of the epileptiform activity (which occurred 3 minutes after the application of BIC) the frequency of epileptiform activities rapidly increased over each subsequent minute (Fig. 5C). Six minutes after the BIC application, spatial mapping of large voltage deflections (10-59 Hz) through RMS detections revealed these clear and repetitive voltage deflections (likely correlated to epileptiform activity) within a few millimeters around the BIC application point during the case.

**Figure 5.**
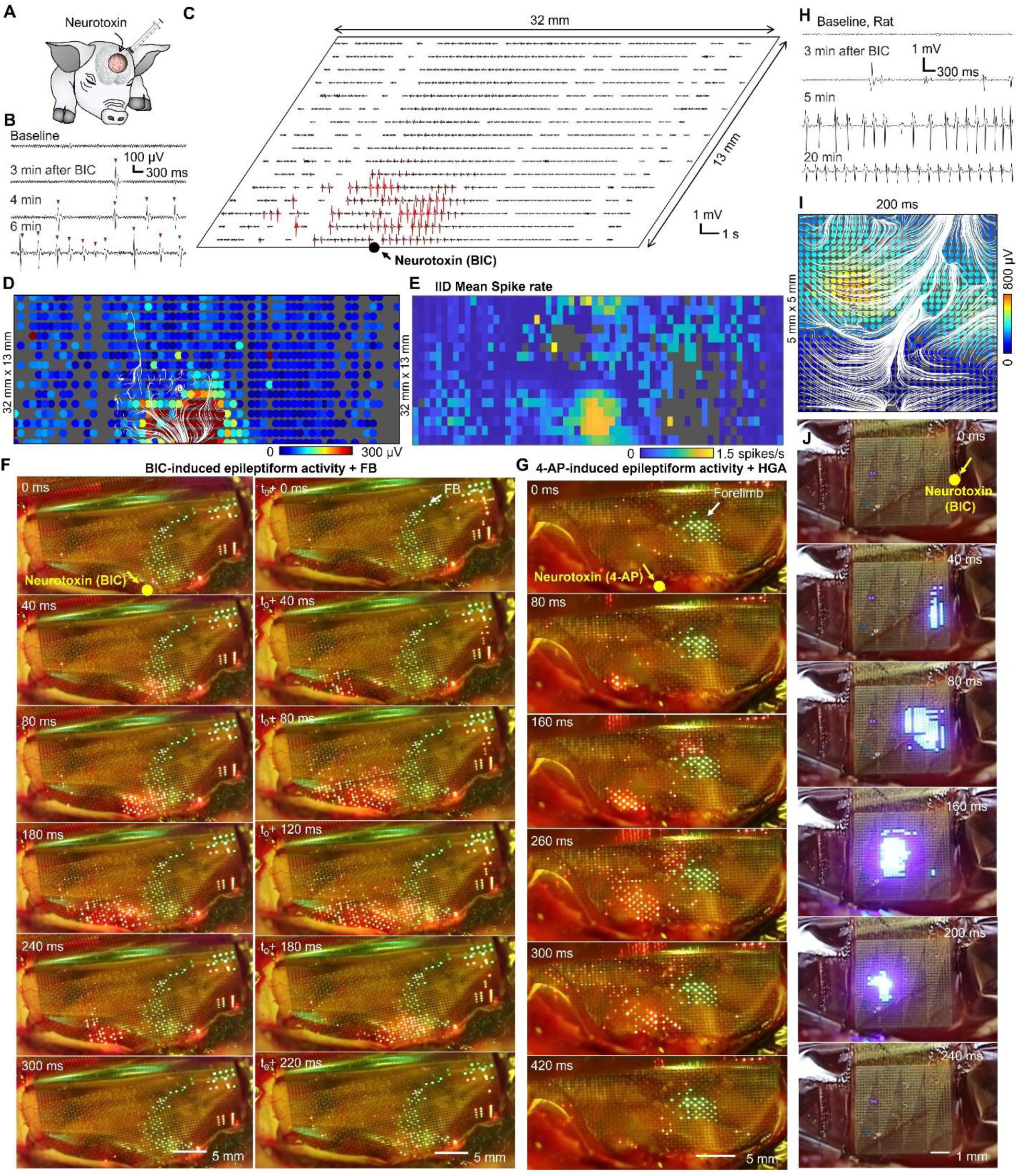
Brain-iEEG-Microdisplay project near real-time display of epileptiform activity from the cortical surface. (A) Schematics of experimental setup for evoking epileptiform activities on the pig brain by epileptogenic neurotoxin including bicuculline (BIC), 4-aminopyrodine (4-AP), and penicillin G (Pen-G). (B) Spatial mapping of waveforms showing locally induced epileptiform activities around the topical application point of BIC crystals. (C) ECoG waveforms of baseline activity before the BIC, and 3, 4, and 6 min after the application of BIC. Spatial potential mapping of epileptiform spike overlaid with white streamlines depicting the propagating direction of the brain wave at (D) 0 and (E) 540 ms. (F) Series of potential maps displayed on the cortical surface (under 550 nm blue blocking lens) with static ‘green’ color showing the motor/sensory functional boundary and dynamic ‘red’ color showing the online potential map of putative epileptiform activity under the BIC application on the cortex. BIC application point is marked as a yellow dot in the figure. (G) 4-AP induced putative epileptiform activities on a different pig with green spot indicating the cortical area responsive to forelimb stimulation as identified with HGA mapping. (H) Waveforms of baseline activity and epileptiform activity induced on the rat brain by BIC over 3, 5, and 20 min. (I) Spatial mapping of putative IIDs together with the white streamlines showing the propagating direction of the putative epileptiform activities. (J) Displaying the putative epileptiform activities induced by BIC application. The original BIC application point is marked as a yellow dot.

We conducted further investigation into the origin and spatiotemporal dynamics of epileptiform activity using streamlines to indicate the propagating direction of brain waves.(*21*) The spatial mapping of epileptiform discharge events was analyzed using the spatial mapping of potentials in band-pass filtered data (10-59 Hz) and streamlines derived from phase gradient analysis.(*42*) The epileptiform activity (Fig. 5D) revealed an epileptiform discharge wave originating near the BIC application site which then propagated outward as represented by the white streamlines (Fig. 5D). Conversely, the second epileptiform event occurred about 500ms after the first and showed the epileptiform discharge wave propagating in the reverse direction, back toward the origin of the first epileptiform discharge. To ascertain whether the observed epileptiform waves were pathological interictal discharges (IID), we utilized offline automatic detection algorithms commonly employed for identifying IIDs(*43*) with visual confirmation by an experienced epileptologist (SSC). We found overlap between the detected IIDs and the spatially located RMS detections near the site of the BIC application (Figs. 5C,D,E). This spatial mapping of the spike rate for the detected voltage deflections possibly correlating to IIDs (Fig. 5E) correlated to the displayed RMS-measured voltage changes on the surface of the pig’s brain using the Brain-iEEG-microdisplay (Fig. 5F). The images were captured under blue-light blocking glass (>550 nm long pass filter, Fig. S2) to enhance the distinction between the green and red colors displayed by the dual-color μLED arrays. While displaying the M1/S1 boundary as a static green line, the large voltage deflections possibly correlating with IIDs were shown in near real-time in red. Using this Brain-iEEG-microdisplay, a pathological wave likely correlating to an IID was observed originating near the BIC application site which propagated to the left side. A consecutive event then showed these voltage deflections propagating back in the reverse direction (Fig. 5F, Movie S3). These observations are consistent with the findings from our offline analysis (Figs. 5D,E). The voltage waves likely correlating with IIDs propagated back and forth repeatedly from the BIC application site throughout the entire one-hour experiment duration. Moreover, when another BIC dose was administered on the rightmost side of the grid, a large voltage wave originating and propagating from the new application site was observed (Fig. S8). Similar propagating behavior of pathological waves was observed when 4-AP (Fig. 5G) or Pen-G (Fig. S9) were injected sub-cortically.

We expected the clinical application of the Brain-iEEG-microdisplay to detect and display pathological activities which would be co-registered to functional regions in the cortex. To this end, the cortical regions responding to electrical stimulation of the forelimb were mapped and displayed as a static green spot while near real-time evolution of the pathological activity was shown dynamically as it occurred in red (Fig. 5G) (N=2 pig cases). The large repetitive voltage waveforms which are likely IIDs originated near the 4-AP injection site and propagated and spread away from that site. This shows that the Brain-iEEG-microdisplay technology could be used to communicate multiple levels of information, display functional boundaries in green color, and show high-resolution near real-time videos of pathological waves with red.

Lastly, we evaluated the Brain-iEEG-microdisplay’s capacity to detect and display fine scale pathological activity at a high-resolution. The Brain-iEEG-microdisplay was positioned on the barrel cortex of a rat after which epileptiform activity was induced by administering BIC. Similar to the case of the pig, the first voltage event which are putative IID occurred 3 minutes after administering BIC to the barrel cortex of the subject rat. The frequency of these voltage deflections increased to 4 spikes per minute at the 5-minute mark (Fig. 5H), with a substantial peak-to-peak amplitude of 3 mV. After 20 minutes had elapsed since the first BIC dose, the voltage deflections amplitude decreased to 1 mV peak-to-peak, and the same spiking rate of 4 spikes per minute continued from the 5-minute mark. We also applied propagating wave analysis to the observed events by overlaying the potential map (filtered in the frequency range of 10-59 Hz) and streamlines representing the instantaneous propagating characteristics of pathological brain waves analyzed by the phase gradient (Fig. 5I). These putative IIDs on the rat brain surface were visible in near real-time, as shown by the series of images at each time point (Fig. 5J) and supplementary video (Movie S4). We repeatedly observed the IIDs originating near the BIC application site and propagating and spreading out to the left side. These findings demonstrate that the Brain-iEEG-microdisplay is capable of displaying submillimeter-scale pathological wave dynamics in near real-time.

## DISCUSSION

The Brain-iEEG-microdisplay displayed neurodynamics of brain cortical activity in near real-time and enabled high-resolution mapping of pathological activities in sub-mm resolution over the extent of centimeter range. The close proximity of the µLED system and µECoG grid led to interference, causing high-frequency noise to be added to the ECoG signals when the µLED system was powered on. This interference resulted from the microdisplay being driven by high-frequency 5V pulses, even when the LEDs were not emitting light. The noise appeared as multiple peaks in the power spectral density plots, typically starting at 98.63 Hz and its harmonics (see Fig. S10). Despite these noise issues, we were still able to effectively extract meaningful neurophysiological signals, displaying important cortical functional boundaries and propagation of IIDs. However, by utilizing customized LED driving hardware and by invoking ground planes on the flexible connectorization of the device, we believe that the noise caused by the interference could be further reduced. To translate this technology to human application, the system must be able to perform continuous and higher sensitivity measurements for the detection of leakage currents preferably on each metal line. This approach could replace the intermittent impedance measurements performed every 10 s. While the PtNRGrid passed the biocompatibility, sterility, and packaging requirements for clinical translation, these tests must be performed on the completely assembled Brain-iEEG-microdisplay for use in humans. For full color display, lamination of three red, green, and blue GaN μLEDs will be required as opposed to the color conversion scheme presented here for binary display of normal and pathological brain activity. With the increase of the spatial resolution toward high-definition display, the transparency of the Brain-iEEG-microdisplay can be compromised. However, with the high-frame display, it is possible to project the anatomical features as a static image on the Brain-iEEG-microdisplay and simultaneously display normal and pathological activity in near real-time. Lastly, it would be desirable to excise tissue while the Brain-iEEG-microdisplay is placed on the brain’s surface without interfering with its function. We envision that additional transducers can be integrated on the PtNRGrid to perform micro-excisions for diseased tissue without compromising its conformity. With light directed toward the cortical surface, the Brain-iEEG-microdisplay can be used for high-resolution optogenetic stimulation.

We have developed and demonstrated the use of the Brain-iEEG-microdisplay for direct display of cerebral cortical activity. The Brain-iEEG-microdisplay is composed of flexible 1024 or 2048 µLED arrays, combined with 1024 channel PtNRGrids and associated hardware and software for near real-time display. With its high spatial and temporal resolution of neural activity and near real-time visual feedback, the Brain-iEEG-microdisplay offers a significant improvement over current intraoperative brain mapping practices. The flexible µLED arrays built on thin, semi-transparent, and conformal polyimide substrates can be reconfigured in pitch to achieve significant cortical coverage. Multiple colors can be displayed to indicate different types of cortical activity. The Brain-iEEG-microdisplay is scalable, has the potential to improve neurosurgical mapping strategies by providing near real-time feedback on neural activities in the treatment of brain disorders. This could lead to better patient outcomes and could also be used to further neuroscientific investigations.

## MATERIALS AND METHODS

Experimental methods are enclosed in detail in the Supporting Information.

**Supplementary Materials**

Materials and Methods

Figure S1-10

Tables S1-S2

Movie 1-4

## Acknowledgements

The authors are grateful for the technical support from the nano3 cleanroom facilities at UC San Diego’s Qualcomm Institute where the fabrication of the Brain-iEEG-microdisplay was conducted. The authors are thankful to Qromis, Inc., for providing the GaN LED substrates. The authors are also grateful for the staff support at the Center of the Future of Surgery at UC San Diego where the pig animal surgeries and experiments were conducted. This work was performed in part at the San Diego Nanotechnology Infrastructure (SDNI) of UC San Diego, a member of the National Nanotechnology Coordinated Infrastructure, which is supported by the National Science Foundation (Grant ECCS1542148).

## Authors contributions

S.A.D., J.C.Y., A.C.P., and S.S.C. conceived the project. S.A.D. led the project. Y.T. and T.W. fabricated the Brain-iEEG-Microdisplay, and Y. T. designed the tasks, developed the software, and conducted all data analysis with S.A.D.’s guidance and with input from A.C.P, E.H., and S.S.C. P.C.C. contributed to the fabrication of the GaN μLEDs. H.S.U. performed all surgeries. D.M.R. devised the anesthetic sequence and D.M.R. and P.P. performed anesthesia and mentoring in the pig’s operating room. D.K. and A.M.B. contributed to the μLED recording hardware with input from I.G. J.L., D.R.C., and K.L. contributed to the animal experiments. K.J.T. performed histology. The animal experimentation paradigm was developed by Y.T., B.C., J.C.Y., A.C.P., S.S.C., and S.A.D. S.A.D. and Y.T. wrote the manuscript and all authors discussed the results and contributed to the manuscript writing.

## Data and materials availability

All data is in the paper and supplementary materials are available by contacting the corresponding author with reasonable requests while honoring institutional and funding agency guidelines.

