## Supplementary Materials for "The Brain Electroencephalogram Microdisplay for Precision Neurosurgery"

**This file includes Supplementary Methods, Supplementary Figures, and Supplementary References.**

### 1. Fabrication of Brain EEG-Microdisplay

#### 1.1. Fabrication of Flexible, Multi-Color Micro-LED Arrays

A monolithic, scalable process was developed for the fabrication of flexible GaN micro-LED arrays on 6-inch engineered substrate. The process leveraged on the low-defect density GaN epitaxially grown on a newly commercialized Qromis® Substrate Technology™ (QST) based on polycrystalline AlN (poly-AlN) with matched thermal expansion coefficient (GaN-on-QST) (Fig.

S1a). An 10-µm-thick GaN epitaxial layer grown on Si(111)/$SiO_2$/poly-AlN consisted of *p*-type GaN (150 nm), InGaN/GaN multiple quantum wells, and *n*-type GaN layers (Fig. S1b).

Micro-LED die was designed to have circular shape to minimize the sharp corners to minimize the potential mechanical damage on the biological tissues. To build a working LED, ohmic contacts to p- and n-type GaN layers were first formed. *p*-type ohmic contact was prepared by treating the surface of *p*-type GaN with aqua regia for 5 min and consecutively depositing Cr/Au (10/10 nm) layers [Ref: Cr/Au ohmic contact with p-GaN]. To expose the n-type GaN, GaN layer was etched through $BCl_3$/$Cl_2$ reactive ion etching (RIE) (Oxford Plasmalab80) to expose n-type GaN layer. The exposed n-type GaN was cleaned by diluted HCl and Cr/Au (10/100 nm) contacts were selectively deposited on n-type GaN layer. Individual dies were defined by doing 10 µm deep isolation etching through a thick $SiO_2$ mask (4 µm) using inductively coupled plasma reactive ion etcher (ICP-RIE, Trion). Circular array of $SiO_2$ masking layer were formed by plasma enhanced chemical vapor deposition (PECVD, Oxford) followed by ICP-RIE patterning of $SiO_2$ through the Cr mask (200 nm).

A 5µm polyimide layer (HD2611, HD Microsystems) was coated on the substrate through 1h 350°C curing process in a Carbolite Oven. Via holes to the LED contacts and perfusion holes were patterned (MLA150, Heidelberg) and etched (ICP-RIE, Trion) by a 100nm sputtered Ti mask (Denton Discovery 18). Then we sputtered Cr/Au (50/500nm) metal leads on this polyimide layer to form the connection to all p contacts of LEDs. A similar round of process was done including second polyimide layer deposition, via holes to the 1$^{st}$ metal leads and perfusion holes etching, and the metal leads deposition connecting all n contacts of LEDs. A third polyimide layer was deposited to passivate the metal leads, followed by via holes etching to expose the metal connection area. Different devices on the substrate were lifted-off in Buffered Oxide Etchant solution in two days. After lift-off process the Si layer on the back side of individual device was removed through $XeF_2$ Etcher (Xactix).

1.2. Fabrication of PtNRGrids

The fabrication process of Brain EEG-Microdisplay involved creating PtNRGrids with 1024 channels that were specifically designed to match the dimensions of micro-LED arrays. The process starts with using a polished and cleaned soda lime glass plate with dimensions of 7” × 7” × 0.06” as the substrate. A 3.7-µm-thick-parylene C layer is then coated on the glass substrate using a parylene deposition system (Specialty Coating Systems 2010 Labcoter), and metal leads are formed on the parylene C layers by lithography, descum, metal deposition, and lift-off process using micro-fabrication tools such as maskless photolithography system (Heidelberg MLA150), UV flood exposure system (DYMAX), plasma etcher (Oxford Plasmalab80), and e-beam evaporator (Temescal). The metal leads are composed of Cr/Au/Cr/Au (10/250/10/250 nm) formed by two cycles of patterning, deposition, and lift-off steps. A PtAg alloy is formed on individual recording sites by photolithography (Heidelberg MLA150), descum (Oxford Plasmalab80), and PtAg alloy co-sputtering (Denton Discovery 18). A Ti capping layer is deposited on top of PtAg alloys to prevent oxidation during the following processes. After depositing the second parylene C layer (3 µm) to conformally cover all the metal leads and PtAg alloys, via holes and electrode outline etching were performed through a patterned Ti hard mask using a reactive ion etching process (Oxford Plasmalab80). The electrodes are lifted-off from the substrate and de-alloyed on the surface of 60°C nitric acid to form the non-toxic platinum nanorods (PtNRs) with low electrochemical impedance. More detailed fabrication processes and characteristics of PtNRGrids could be found elsewhere.(*1*)

1.3. Bonding and Assembly of Brain EEG-Microdisplay

The flexible μLED was connector to the LED driver electronics through a flexible printed circuit board (FPCB) and connectors. A selectively-deposited silver epoxy (8331, MG Chemicals)-based bump bonding was used to make reliable bonding interface between the thin μLED layer and customized FPCB. For the multicolor applications, red and green indium phosphide/zinc sulfide (InP/ZnS) quantum dot color conversion (QDCC) inks (Mesolight) were printed on top of the GaN dies through ink-jet material printer (Fujifilm Dimatrix 2850). InP/ZnS quantum dots were selected because they have lower potential toxicity compared to other types of quantum dots. [Ref: Brunetti, Virgilio, et al. "InP/ZnS as a safer alternative to CdSe/ZnS core/shell quantum dots: in vitro and in vivo toxicity assessment." Nanoscale 5.1 (2013): 307-317.] A 50-nm-thick $Al_2O_3$ passivation layer was deposited on the QDCC-printed devices using atomic layer deposition (Beneq TFS200) to minimize the moisture or oxygen exposure of the QDCC ink that may degrade the quantum efficiency of QDCC ink over time.(*2*) A 3.3-μm-thick Parylene C layer was then coated (Specialty Coating Systems 2010 Labcoter) to further passivate the device to be used in wet environment such as brain surface. The QDCC ink was then photo-activated by UV exposure under an inert nitrogen environment to enhance the luminescence efficiency.(*3*)

To bond the PtNRGrids to the extender PCB, a selectively-applied silver epoxy (MG Chemicals 8331) was used on the PCB footprints. A stencil mask made of silicone adhesive PET tape (Advanced Polymer Tape) with laser-cut holes was used to apply the silver epoxy selectively onto the PCB footprints. The PtNRGrid was temporarily placed on a clear plastic film (Steriking®)/glass plate and micro-aligned with and pressed against the extender PCB with silver epoxy bonding bumps. They were then cured on a 75°C hotplate for 15 minutes under 5-10 N force to ensure complete curing of the silver epoxy and establish electrical connections across all bonding contacts. The electrically bonded PtNRGrid was released from the glass plate, and the electrodes were characterized on a benchtop. Before bonding, the PCB edge was ground and smoothed to minimize potential damage to the thin metal leads in parylene C films.

To create the Brain EEG-Microdisplay, the PtNRGrids were assembled with a flexible μLED array. Two separate grids with matching dimensions were precisely aligned using micro-alignment stages with four-axis degrees of freedom. Once the individual μLED dies and PtNR recording sites were aligned, a small amount of photoresist (AZ5214E-IR) was applied to the 'neck' part of the μLED array and PtNRGrids through holes prepared in the μLED array. This held the two thin and flexible grids together (Fig. 1c) with an added feature that the μLED layer could be lifted-off from the PtNRGrids when needed. For the 32mm x 32mm Brain EEG-Microdisplay, the perfusion hole arrays in the PtNRGrids were aligned with the through hole arrays in the flexible μLED array. The through hole arrays (0.5 mm diameter, 1 mm pitch) in the Brain EEG-Microdisplay enabled Ojemann probe to directly stimulate the cortical surface through the grid.

### 2. Electronics and Software to Drive Brain EEG-Microdisplay

#### 2.1. Hardware to Operate Brain EEG-Microdisplay

We developed the ORB1024_V1, an acquisition board that utilizes Intan Technologies' RHD2164 chipset, to record from up to 1024 channels concurrently, with a maximum of 30 kilo-samples per second (ksps) per channel. The board consists of 16 RHD2164 chips to amplify and digitize the brain signals, LGA1155 CPU socket that connects to our electrode arrays, and Omnetics 12-pin connectors that interface with an off-the-shelf 1024-channel Intan RHD recording controller. With the ORB1024_V1, we can connect our flexible electrodes array and digitize the 1024-channel analog brain signals robustly. Using the Intan RHD recording controller, we set the sampling rate of the RHD2164 chips as 20 ksps and recorded the ECoG signals with a data rate of around 2.4 GB/min. The recorded streaming data is transmitted from the recording controller to a host computer via the USB interface.

Fig. S5(A) depicts the μLED + μECoG feedback system, which comprises the recording station mentioned above, a host computer, and μLED subsystem with a mechanical toggle switch and an external impedance analyzer. The recording station records neural activity and feeds it to the μLED subsystem to map the neural activity onto the μLED array. As shown in Fig. S5(B) and S5(C), the LED driver board has four LED driver chips (IS31FL3741A, Lumissil) that can support driving the 32 × 32 single-color GaN μLED array with a voltage level of 5 V and an average current of 3.75 mA. Also, the board includes the MCU, Teensy 4.1, to configure the LED driver chips through the 1 MHz I2C interface and to supply power through the USB interface. For the dual-color Brain-iEEG-microdisplay, two LED driver boards were used and controlled individually via the host computer.

Our μLED subsystem supports three operation modes: normal mode, impedance measurement mode, and hardware shutdown mode. In normal mode, the host computer receives 20 kSps recorded brain waves through USB_0, and the Qt software processes the data and sends a series of control bits to the μLED subsystem through USB_1. The MCU transforms the USB-formatted data into the I2C format to program the LED driver chips, resulting in a change of the LED pattern. Through an iterative process of updating the LED driver chips based on recorded brain waves, the μLED subsystem can provide video-rate visualization. To ensure the safety of the brain and the surrounding tissue, the μLED subsystem switches to impedance measurement mode every 10 seconds in normal mode by setting MODE_SEL low. In impedance measurement mode, the μLED subsystem keeps track of the leakage current on the brain and the tissue by measuring the impedance between the μLED array and a reference node. As the reference node, a needle is connected with the ground of the electronic system and inserted into the tissue [See Fig. S5(C)]. Upon entering impedance measurement mode, the μLED driver board electrically ties up all the column and row control lines of the LED matrix into a single node by changing the active channel of the FET multiplexers. After the multiplexers are set, the impedance analyzer inputs the current over the LED array and measures the impedance between the LED matrix and the reference needle. In this work, additional single Intan chipset and its controller are used as an external impedance analyzer. If the measured impedance at 1 kHz is higher than 100 kOhm, the system goes back to normal mode by setting MODE_SEL high. However, if the measured impedance gets lower than 100 kOhm, the μLED subsystem enters shutdown mode, which power down the μLED driver chips, to prevent damage to the brain. The system also includes a manual toggle switch that enables to enter shutdown mode in unexpected situations. The overall flow chart of the μLED subsystem operations is shown in Fig. 7S(D).

### 2.2. Near real-time Data Processing Software of Brain EEG-Microdisplay

The brain data recorded with the 1024-channel PtNRGrids were processed in near real-time to instantaneously display the cortical activity on the microdisplay. This near real-time data processing method was used to display the epileptiform activities (Fig. 5) and extent of electrical stimulation (Fig. 4) from the surface of the brain. To achieve this, the Intan Technologies' C++/Qt open source RHD Recording Controller software (Version 2.08) was customized to perform high and low pass filtering and spatial mapping of RMS potentials on all channels. The computer received 25 ms of data packets containing 1024 channels from Intan recording controller. The 25 ms data packet was bandpass filtered in 10-59 Hz frequency window, and root-mean-square (RMS) value of filtered data was calculated for individual channels to generate a single frame of spatial map of the RMS potentials. These RMS mapping data were displayed on the processing computer screen, and serial communication was used to send the spatial mapping information to the microdisplay via a Teensy 4.1 microcontroller. The resulting refresh rate of the microdisplay was 40 Hz.

Trial averaging was used to show the M1/S1 boundary and the localized high gamma activities of the animal under repeated electrical or mechanical sensory inputs. A sensory input to the animal was configured to send a time-locked TTL signal to the Intan recording controller. Once a TTL signal was captured, an evoked response waveform typically between 0 to 100 ms post-stimulus was stored in the memory which waveforms were updated by trial averaging with the additional inputs of evoked responses. The typical post-stimulus time range used for RMS mapping in the Brain EEG-Microdisplay was configured to be between 18 and 22 ms. The software had a user-input feature that allowed the user to select the post-stimulus time range for calculating the RMS mapping.

To ensure a seamless image on the microdisplay, impedance measurements were performed to screen out channels with high impedance (>100 kΩ at 1 kHz) that showed poor electrochemical impedance, causing high-amplitude noise and large RMS potential. This was necessary to avoid distortion of the actual brain activities. The RMS potential values of the nearest neighboring channels with good impedance (<100 kΩ at 1 kHz) were then used to fill in the RMS potential value on the high-impedance channels, enabling a continuous display of the cortical activity on the microdisplay.

2.3. Noise contribution of LED to the ECoG recording

The close proximity of the μLED system and μECoG grid caused interference between the two systems, resulting in high-frequency noise being added to the ECoG signals once the μLED system was powered on. This interference was due to the microdisplay being driven by high-frequency 5V square pulses; even when all the LEDs are not emitting light, the LED driver (Lumissil IS31FL3741A) sends 5V pulses sequentially to all the rows (*n*-contact) and columns (*p*-contact), setting the net bias voltage to 0. The noise power spectral density plots before and after turning on the LED system (Fig. S10) show multiple discrete peaks starting from 98.63 Hz and their harmonics at higher frequencies. For the near real-time display of cortical activities, we applied a band pass filtering below 90 Hz (typically between 10 and 59 Hz) to the ECoG signals which effectively filtered out the artifactual noise caused by the LED system.

For the trial averaged high gamma activities, although the selected frequency window of 70-190 Hz included some noise peaks from the LED, this noisy signal averaged out with increasing number of trials. To achieve a lower background noise level during the HGA mapping, another strategy was to use a switch that could power on and off all the LED driving chips. The switch was used to turn off the LEDs while recording the brain activity in response to the sensory tasks. The recorded responses were averaged over multiple trials. After the sensory task was completed, the high gamma activity (HGA) map was displayed by turning on the LEDs.

2.4. Offline Data Analysis

Together with the near real-time processing to display cortical activities on the brain, the ECoG signals from 1024 channels were recorded for offline analysis. Individual recording channels were mapped to the individual spatial coordinates on the PtNRGrids, and this mapping was tabulated in a spreadsheet for each electrode type. This allowed us to spatially display waveforms, RMS potentials, and impedance magnitude using the recorded data. To ensure accuracy of the analysis, channels with an *in vivo* impedance magnitude above 100 kΩ at 1 kHz were excluded from analysis, as they usually showed a response that was artificially large and were more susceptible to noise. Additionally, neighboring channels with very low impedance magnitude (< 1 kΩ at 1 kHz) were evaluated as potentially shorted channels and were excluded from the offline analysis. Furthermore, all recorded signals (unless explicitly specified as "raw") underwent processing to eliminate 60 Hz and their noise harmonics using digital notch filters.

For the M1/S1 sensory boundary localization, signals were digitally filtered in the frequency window of 10-3000 Hz using a Butterworth 4th order filter with MATLAB's "filtfilt" function. No re-referencing was used since this could potentially cause an undesirable offset in SSEPs. TTL pulses time-locked to the electrical stimulation of forelimb was used to determine time epochs for trial averaging (N=50) the SSEPs.

The HGA mapping was carried out by trial averaging the raw waveforms based on the TTL pulses time-locked to the air-puff or electrical stimulation. We then re-referenced the recorded signals by subtracting the common-averaged signal across channels. The common-average was calculated either by taking the average of all the working channels or by taking the average of a few selected channels.(*1*) This effectively removed the motional artifacts, electrocardiogram, and low frequency noise. The signals were digitally filtered using a Butterworth 4th order filter under selected frequency windows of 70-190 Hz, and 50 trials were aligned and averaged based on the TTL pulses that triggered air-puff or electrical stimulation. All digital filters were implemented in Matlab using the zero-phase distortion filtering function, "filtfilt", which effectively doubled the filter order to 8. The amplitude of the signals in each frequency window were calculated by taking root-mean-square (RMS) of the absolute value of the Hilbert transformed signal in a 15~25 ms time window after the stimulation.

The propagating dynamics of the beta waves were calculated by taking the spatial phase gradients of the beta waves following the methods described in Rubino *et al.(4)* and Muller *et al.*.(*5*) The signal was first filtered in beta band of 9-18Hz using a Butterworth 4$^{th}$ order filter with the "filtfilt" function in Matlab. The phase angle of the beta wave for each channel was calculated by taking the inverse tangent of the imaginary part over the real part of Hilbert transformed data, and the phase was unwrapped over time. The propagation directions of the beta waves calculated from the spatial phase gradient were represented as a vector field and streamlines, were used to visualize the long-range propagation directions of the waves. The streamlines were plotted using the streamline function in Matlab with a default setting.

We used offline analyses to detect interictal discharges (IIDs) using an automatic IID detection algorithm (version v21, default settings except -h at 60; http://isarg.fel.cvut.cz) (*6*) The automatic IED detection algorithm adaptively models distributions of signal envelopes to discriminate IIDs from intracranial recordings. (*6*)

### 3. Pig Experiments

#### 3.1 Pig Models and Task Information

Table S1 summarizes the pig models, devices, and task information.

#### 3.2. Anesthesia and Pre-operative Preparation of Pigs

All procedures for the pig experiment were approved by the UCSD Institutional Animal Care and Use Committee under protocol S19030. The pre-operative medication included a cocktail of Ketamine, Midazolam and Atropine - administered IM. An 18G catheter was placed into an ear vein and tracheal intubation was performed with the aid of a laryngoscope. A cuffed endotracheal tube was placed, secured, and connected to an anesthesia machine equipped with a charcoal canister as a scavenger. Mechanical ventilation was initiated with a tidal volume set at approx. 10 cc/kg at a rate of 8-14 breaths per minute and lactated ringers or saline were infused through an intravenous line at 5-10 ml/kg/hr. Isoflurane anesthesia was maintained using 1-3% isoflurane in 100% oxygen. The animal's depth of anesthesia was continuously monitored by observation of vital signs and the animal's reflex responses. Vital signs including HR, RR, $SPO_2$, temperature, and $ETCO_2$ were monitored and recorded every 15 minutes. The monitored reflexes included the palpebral and pedal responses, as well as jaw tone strength. End tidal $CO_2$ ($ETCO_2$) was monitored and maintained between 35-45 mmHg. A fluid warmer and/or Bair Hugger were utilized in all studies to maintain normal body temperature. Clipping of the hair was performed at the surgical sites. Once the brain EEG-microdisplay was placed on the cortex, the animal was transitioned from isoflurane to IV propofol anesthesia for the remainder of the experiment. During the analysis of epileptiform discharges, a paralytic agent (vecuronium) was readily available in case of a severe motor seizure response in the animal. Only minor motor responses were observed in response to the epileptogenic agents and vecuronium administration was unnecessary. In one animal, vecuronium was administered to reduce shivering artifact that was unresponsive to adequate anesthesia and temperature control.

#### 3.3. Surgical Procedures for Pig Craniotomy and Brain EEG-Microdisplay Implantation

The surgical site was centered over the motor cortex of the frontal lobe and somatosensory cortex of the parietal lobe. Once anesthetized, the animal was mounted into a stereotaxic frame. After immobilization, a skin incision measuring 2-4 inches in length was made. The skull was then exposed using a chisel and retractors, and a bilateral craniotomy was performed using a neurosurgical drill. The skull was removed to create a window approximately 40 mm × 40 mm. The dura underlying the skull was cut open except the dura near the midline. Once exposed, the cortical surface was hydrated throughout the experiment with saline. A brain EEG-microdisplay was then placed on the surface of the exposed cortex.

3.4. Sensory Tasks in Pig Experiments

Electrical stimulation of pig was carried out with 12 mm twisted-pair subdermal needles (Natus) and Intan RHS recording and stimulation system. The subdermal needles were poked into the skin by 12 mm with typically 2 mm separation between the pair of needles. Biphasic current pulses with amplitude of 2 or 10 mA and pulse width of 1.0 ms were delivered to the bipolar needles to stimulate various parts of the pig. The parts we stimulated include forelimbs, snout, cheek, and tongue. Since the maximum current level that could be generated from a single channel in Intan RHS system was around 2.5 mA, we shorted 5 channels and connected them to a single needle and sent time-locked current pulses to 5 channels at once to achieve 10 mA. The other needle was connected to the ground/reference pin in the RHS headstage.

Snout and tongue of pigs were locally stimulated with an air-puff stimulator using the Pneumatic PicoPump (WPI, PV830). Air-puff was delivered through a 1 mm diameter glass microcapillary tube with a pressure of 40 psi. Each air-puff stimulation position was stimulated 50 times, once every 1 s.

Both the air-puff and electrical stimulation were time locked to the recording system by sending TTL signals to both the stimulator and the Intan recording controller.

3.5. Direct Electrical Stimulation of Pig Brain

Pig brain was electrically stimulated at the surface or at depth using clinical stimulators. A Ojemann probe was used to deliver 3 mA biphasic pulses of current at 50 Hz with variable stimulation time ranging from 0.1 to 2 s. Multiple positions on the pig's cortical surface were stimulated either through or around the Brain-iEEG microdisplay. 16 channels stereoelectroencephalography (sEEG) electrode (PMT Corp.) with contact spacing of 6 mm and diameter of 0.8 mm were manually inserted into the depth of the pig brain by 6 cm. Intan RHS system was used to deliver biphasic current pulses of 1~10 mA (0.5 ms positive, 0.5 ms negative) to the adjacent channels on the sEEG electrode. To generate a 10 mA current level, we combined the output of 5 channels in the Intan RHS system by shorting 5 channels to a single needle.

3.6. Epilepsy Pig Model

Epileptic discharges were acutely introduced on the pig brain using epileptic discharge inducing drugs approved under protocol S19030. We used three different epileptic discharge inducing drugs with different application methods on four different pigs including i) topical application of crystals of 1(S),9(R)-(-)-bicuculline methiodide (BIC, Sigma-Aldrich 14343) on the pig's cortical surface (N=1 pig case), or subcortical injection of ii) 10 µL, 100 mM 4-aminopyridine (4-AP, Sigma-Aldrich A78403) solution (N=1 pig case) or iii) 10 µL, 100 mM benzyl-penicillin (Pen-G, Sigma-Aldrich 13752) solution (N=2 pig cases). The drugs were reapplied every 15 min for a total of 3 applications to keep up the efficacy of the effect. All three drugs induced clear epileptic discharges, while the efficacy, amplitude, and spatial distribution of epileptic discharges varied between the three drugs studied. Of the three drugs, BIC produced the highest amplitude epileptiform activity. Notably, controlling the amount of BIC was challenging given its crystalline form which may have led to an overdosing of BIC compared to the other drugs. During our study on anesthetized pigs, we observed mild motor seizure activity near the snout of one pig (Fig. S9), while we did not observe any other motor seizure activity in the other pigs after careful full body observation.

3.7. Histology of the Pig Brain

The brain under the iEEG-microdisplay was resected and immediately immersion fixed in 4% PFA for 24 hours before being transferred to 30% sucrose. The brain was blocked and sectioned at 50 μm on a cryostat and slices were stored in PBS + 0.01% Sodium Azide before further processing. Control and experimental sections from each block were processed in the same well to limit variability. Slices were washed for 2 hours in PBS and blocked for 1 hour in 5% normal goat serum (Jackson Immunoresearch, PA). Sections were incubated with Neurotrace 640/660 (1:50, Thermo Fisher Scientific Cat# N21483) for 30 minutes. Sections were washed for 2 hours in PBS before being mounted and coverslipped with Prolong Gold Antifade Reagent with DAPI. Sections were imaged on a Zeiss Apotome2.

**4. Rat Experiments**

4.1 Rat Models and Task Information

Table S2 summarizes the rat models, devices, and task information.

4.1. Surgical Procedures of Anesthetized Rat Craniotomy

All procedures for the rat experiment were approved by the UCSD Institutional Animal Care and Use Committee under protocol S16020. Male Sprague Dawley rats (10-18 weeks of age) from Charles River were sedated with 3-4% isoflurane and fixed in a stereotaxic frame (Kopf Instruments). Once stable, rats were reduced to 3% isoflurane for maintenance, while monitoring heart rate and blood oxygen levels (Mouse Stat Jr, Kent Scientific). Prior to the craniotomy, contralateral side individual whiskers that were to be stimulated were colored with Sharpies to easily distinguish them in the air-puff stimulation experiment, and the remaining whiskers were trimmed off. A craniotomy was made on the right skull 1 cm lateral and 2 cm posterior from the bregma, exposing the somatosensory (including barrel) cortex, over the right motor cortex. The dura was carefully opened and retracted from the brain, and a small piece of saline-soaked gauze was placed over the brain until the implant was ready. Once the craniotomy was complete, the rat was transitioned from isoflurane to ketamine/xylazine (100 mg/kg ketamine / 10 mg/kg xylazine) and re-dosed every 20-30 min for the duration of the experiment. Temperature, heart rate, and oxygen concentrations were monitored for the entirety of the experiment to ensure adequate anesthesia.

#### 4.2. Sensory Stimulation Tasks on the Rats

The typical size of the craniotomy was $6 \times 6mm^2$, and the $5 \times 5mm^2$ Brain-iEEG microdisplay was implanted covering nearly the entire exposed area of the brain. The reference needle electrode was implanted on the scalp of the rat just next to the craniotomy, and ground was typically connected to a surrounding faraday cage or stereotaxis frame. Individual whiskers were stimulated with an air-puff stimulator using the Pneumatic PicoPump (WPI, PV830). Air-puff was delivered through a 1 mm diameter glass microcapillary tube with a pressure of 20 psi for single whisker stimulation and 40 psi for whole whiskers stimulations. After a 10 s baseline recording, each whisker was stimulated 50 times, once every 1 s. To minimize the chance of stimulating multiple whiskers other than the whisker of interest, whiskers were subsequently trimmed off after each recording. The air-puff stimulations were time locked to the recording system by sending TTL signals to both the stimulator and the Intan recording controller.

#### 4.3. Epilepsy Rat Model

Epileptic discharges were acutely introduced on the rat brain using epileptic discharge inducing drugs approved under protocol S16020. We used two different drugs with different injection methods including topical application of i) crystals of 1(S),9(R)-(-)-bicuculline methiodide (BIC, Sigma-Aldrich 14343) or ii) 50 µL, 100 mM 4-aminopyridine (4-AP, Sigma-Aldrich A78403) solution on the rat's cortical surface just next to the Brain-iEEG microdisplay. The drugs were reapplied every 15 minutes on the same spot to maintain their efficacy, with a maximum of three applications. It is worth noting that when the 4-AP solution was topically applied, it immediately spread over the entire craniotomy, while the BIC crystal in powder form, applied on the cortical surface, slowly dissolved over time on the brain. Both drugs induced clear epileptic discharges, but the response to BIC was more localized on the rat brain. This is likely due to the difference in the degree of spread of the drugs on the cortical surface. During our study on anesthetized rats, we rarely observed motor seizures despite of the huge (> 1 mV) and repetitive epileptiform discharges showing up across the rat brain.

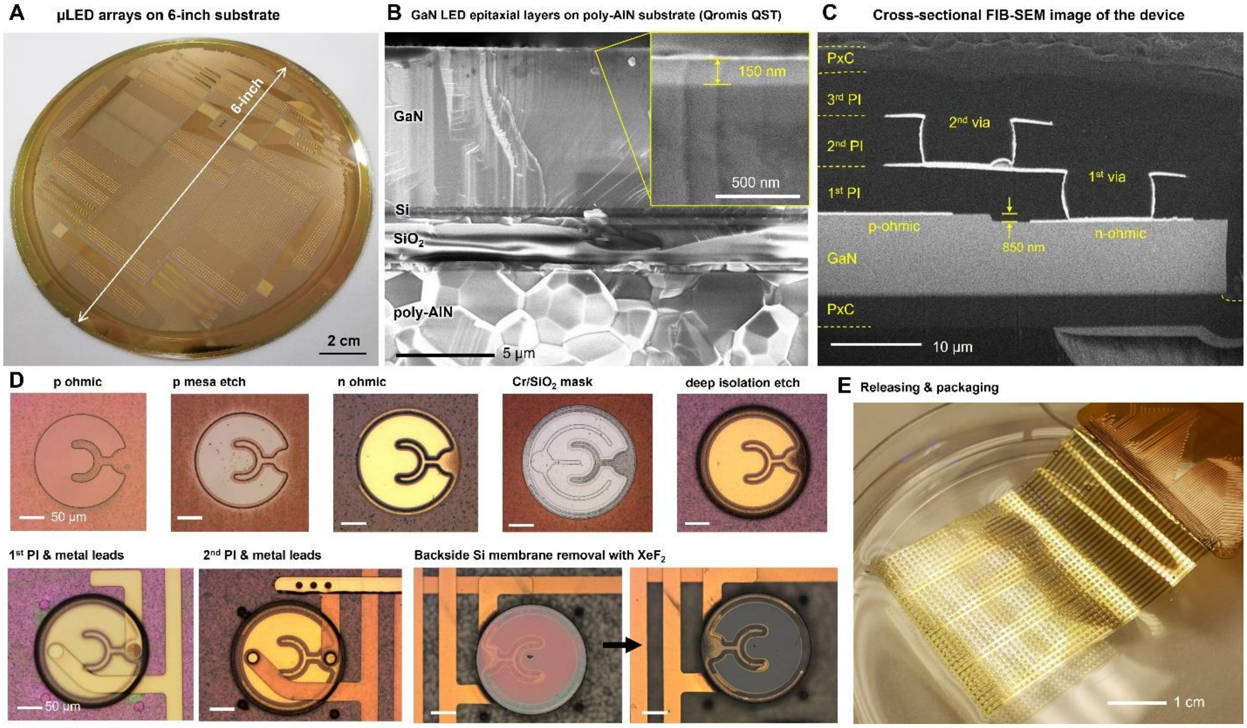


**Figure S1.** Microfabrication procedures of flexible μLED arrays. (A) Photo of μLEDs processed on the Qromis 6-inch QST wafer. (B) Cross-sectional SEM image of QST wafer that consist of poly-AlN substrate, $SiO_2$ intermediate layer, and GaN epitaxial layer with LED structures. 150-nm-thick p-type GaN layer is shown in the subset figure. (C) Cross-sectional FIB-SEM image of the completed μLED with GaN LED die, ohmic contact metal layers, via metallization layers, and multiple polyimide layers and parylene C layers for the crossbar array and electrical passivation. (D) OM image of the microfabrication process including ohmic metal contacts, etching, isolation etching, via metal connection, and backside Si membrane etching. (E) Released flexible μLED layer by dissolving the $SiO_2$ layer shown in (B). μLED is bonded to the FPCB by the silver epoxy bump bonding.

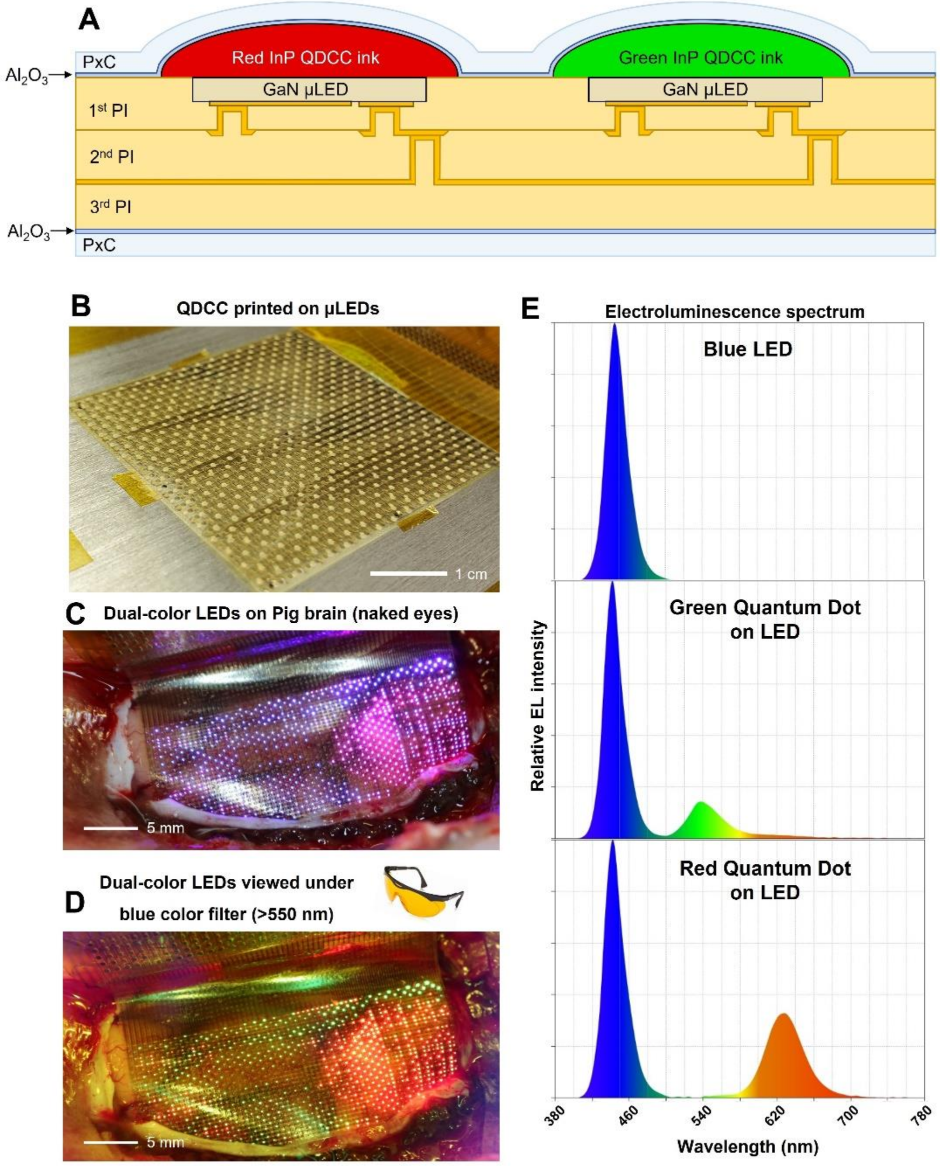


**Figure S2.** Multicolor LED fabrication and luminescence characteristics. (A) Schematic illustration of cross-sectional structure of µLED with red and green InP Quantum Dot Color Conversion (QDCC) inks. Schematics are not drawn to scale. (B) QDCC ink printing on the µLED array by the Dimatix DMP-2850 material printer. Dual-color LEDs on the pig brain showing the motor/sensory boundary (C) under naked eyes and (D) under 550 nm blue light blocking lens. (E) Electroluminescence spectra of blue µLED and color converted µLEDs with red and green QDCC inks.

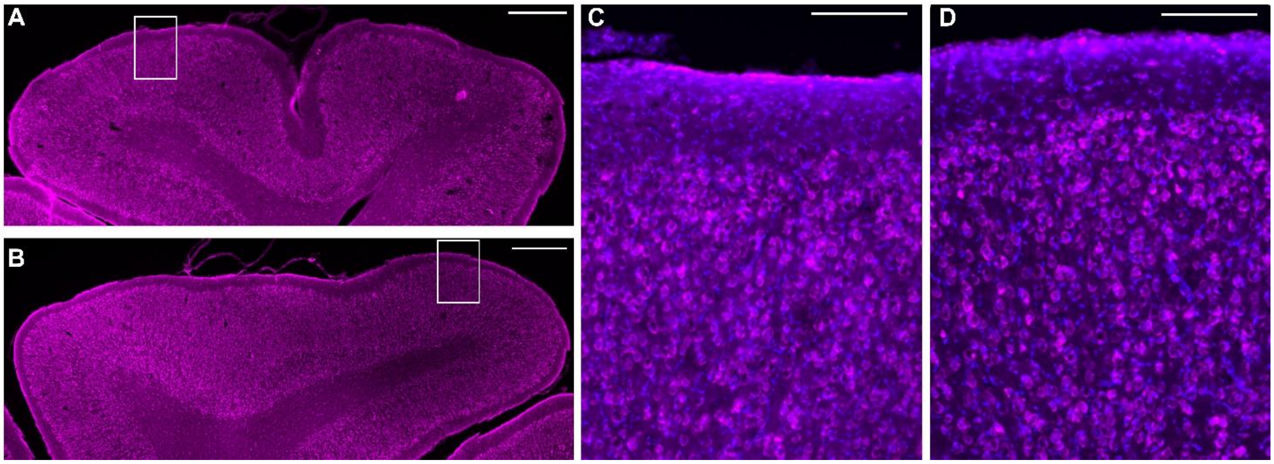


**Figure S3.** Representative images of the cortical surface after Brain-iEEG microdisplay recording. Tiled image of rostrum from the (A) contralateral and (B) grid side with Neurotrace (fluorescent Nissl) staining reveal intact cortical layers with no gross damage on the experimental side (Scale bar = 1 mm). Inset images from the (C) contralateral and (D) experimental demonstrates comparable Layer I thickness, neuron density, and cell morphology (Scale bar = 200 μm).

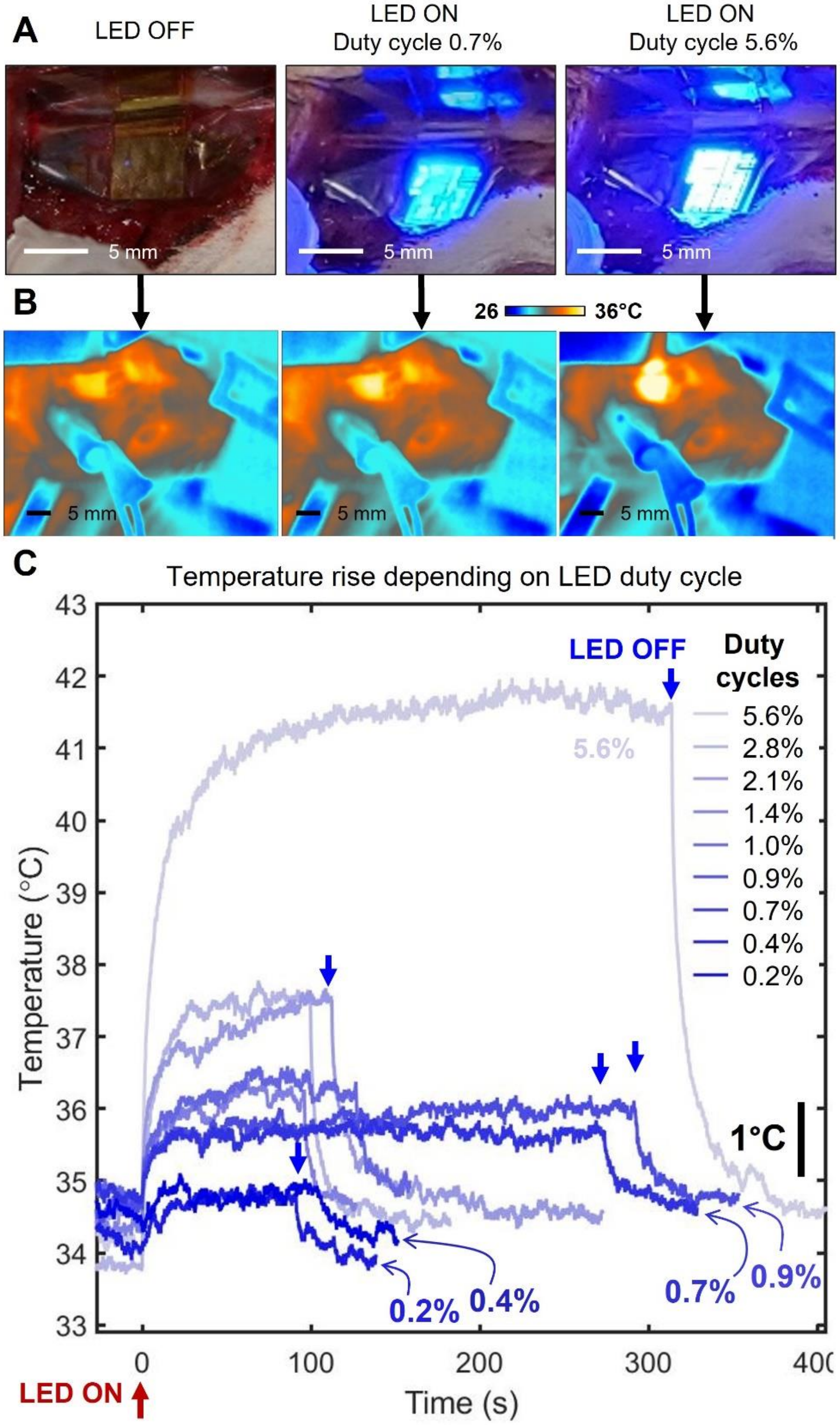


**Figure S4.** Cortical surface temperature of the rat brain under the LED operation. (A) Photo of μLED arrays on the brain with LEDs off and with all the LED pixels turned on at 6 and 50% duty cycles. (B) Corresponding infrared (IR) thermal images when LED was off, or ON at 6 and 50%. (C) Temperature rise and fall with LEDs turned on at different duty cycles and turned off.

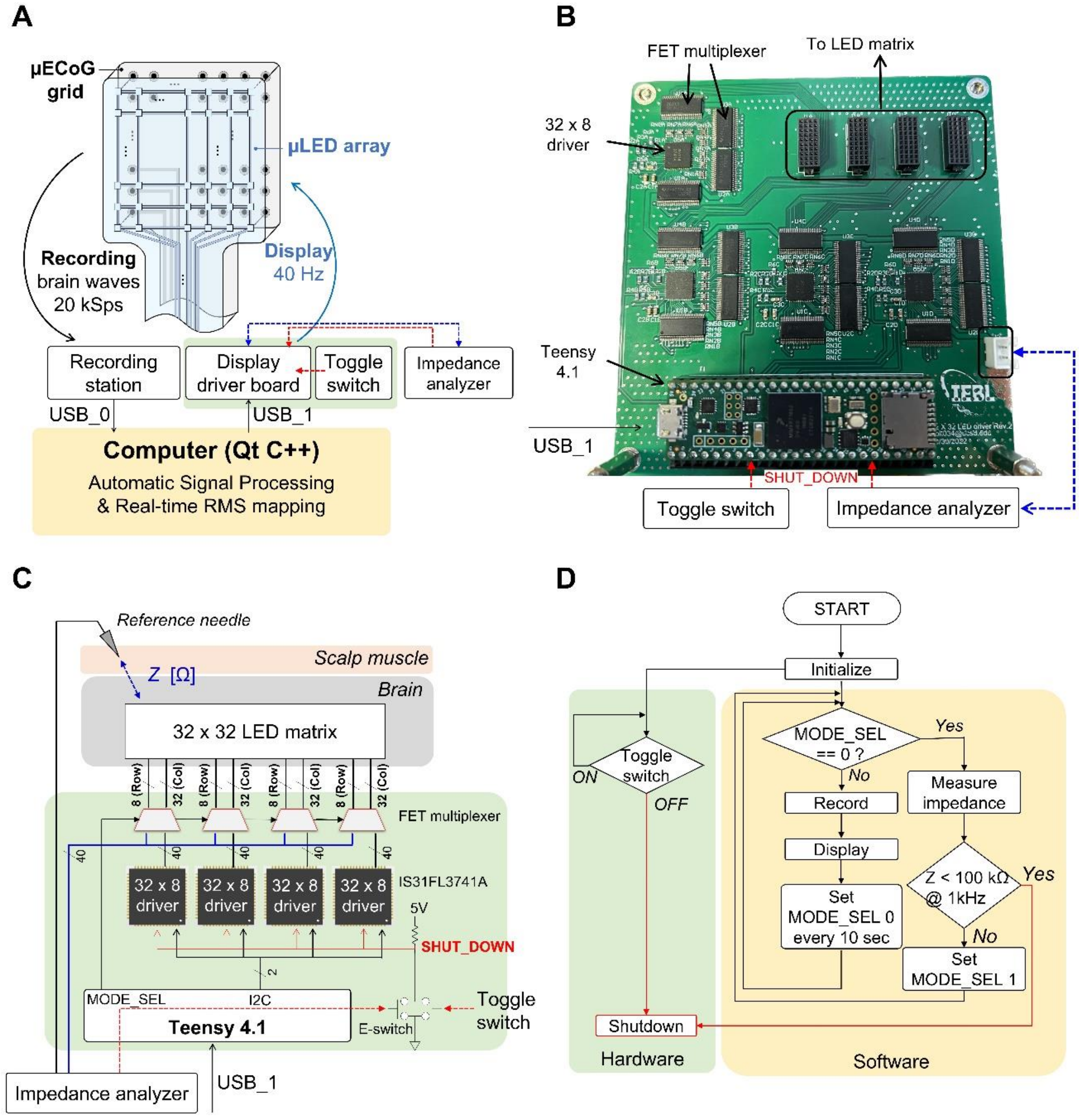


**Figure S5.** μLED + μECoG processing system. (A) Overall block diagram of the feedback system between the μLED and μECoG subsystem, (B) Photograph of the μLED driver board, (C) Schematic of the μLED driver system, and (D) Flow chart of the μLED controlling system.

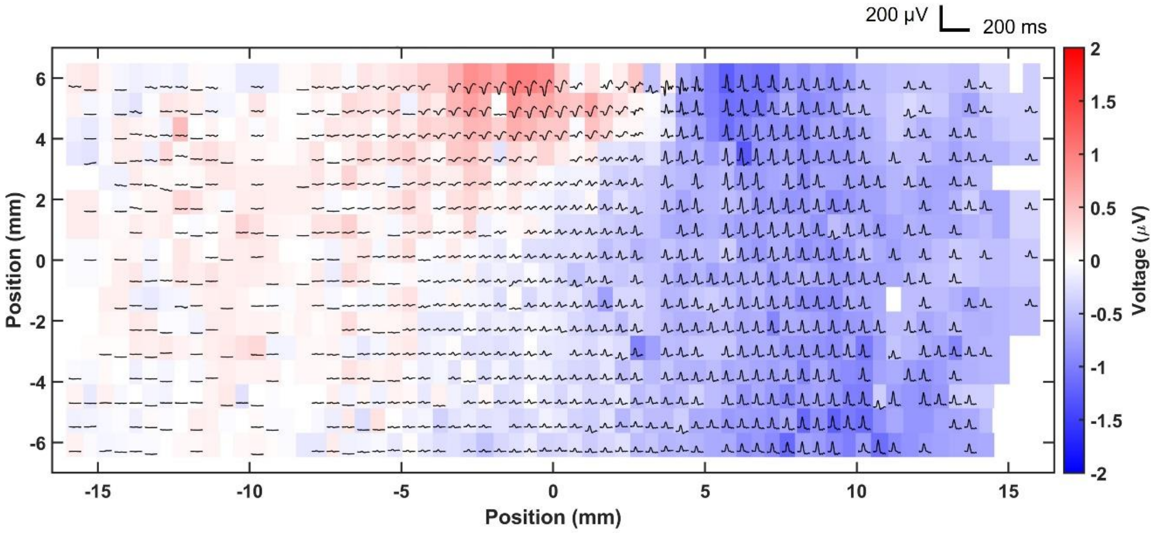


**Figure S6.** Offline analysis of phase reversal mapping of M1/S1 boundary on the pig brain with 13mm x 32mm brain EEG microdisplay device.

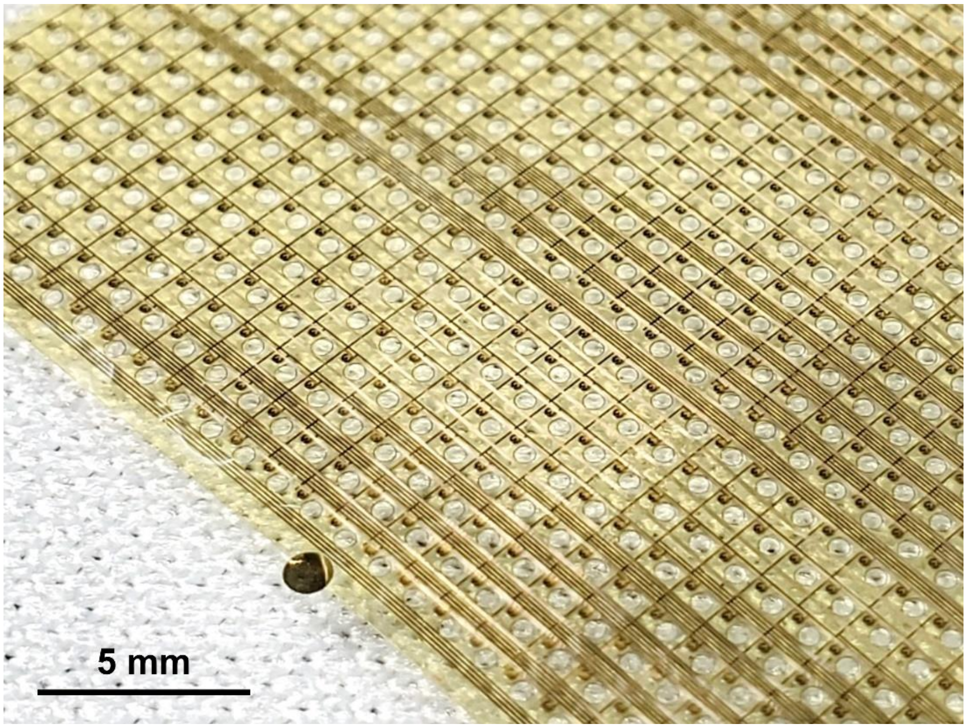


**Figure S7.** Perforated hole array on Brain-iEEG microdisplay that allowed through-grid direct electrical stimulation. The device is placed on a cleanroom woven polyester fabric.

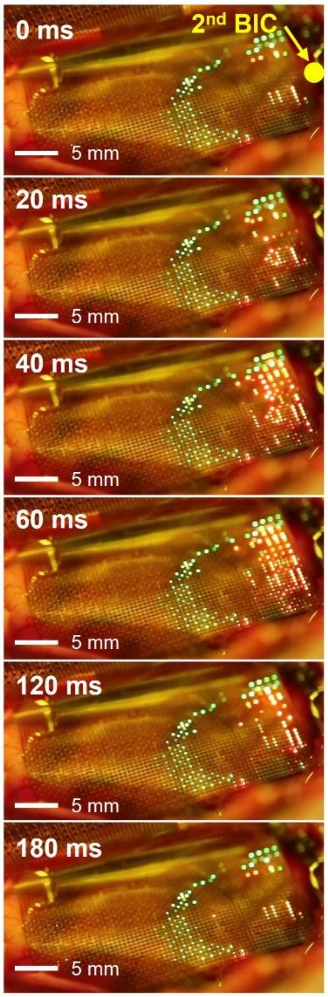


**Figure S8.** Effect of another BIC application on a different spot on the cortex after Figure 5(f). Series of potential maps displayed on the cortical surface (under 550 nm blue blocking lens) with ‘green’ color showing the motor/sensory functional boundary and ‘red’ color showing the online potential map of epileptiform activity under the 2nd BIC application on the cortex. New putative epileptiform activities emerged near the position of 2nd BIC dose and propagated from right to left.

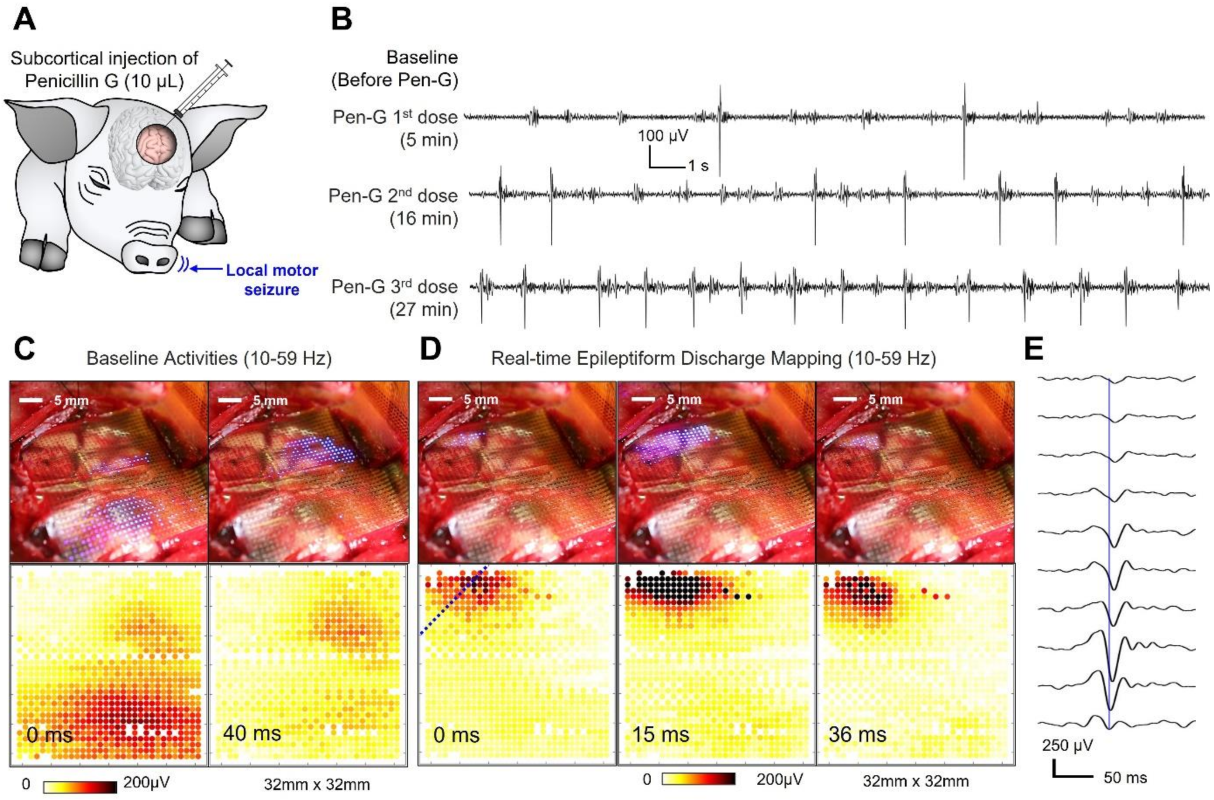


**Figure S9.** penicillin G (Pen-G)-induced putative epileptiform activities display on the cortical surface. (A) Schematics of evoking epileptiform activities on the pig brain by Pen-G). (B) ECoG waveforms of baseline activities before the Pen-G, and 3, 4, and 6 min after the application of Pen-G. (C) Baseline activities of brain waves visualized with LED+ECoG on pig brain and corresponding potential mapping generated by offline analysis. (D) Epileptiform activities of brain waves visualized with LED+ECoG on pig brain and corresponding potential mapping generated by offline RMS potential mapping analysis. (E) Epileptiform spike across the channels along the dotted blue line in (D).

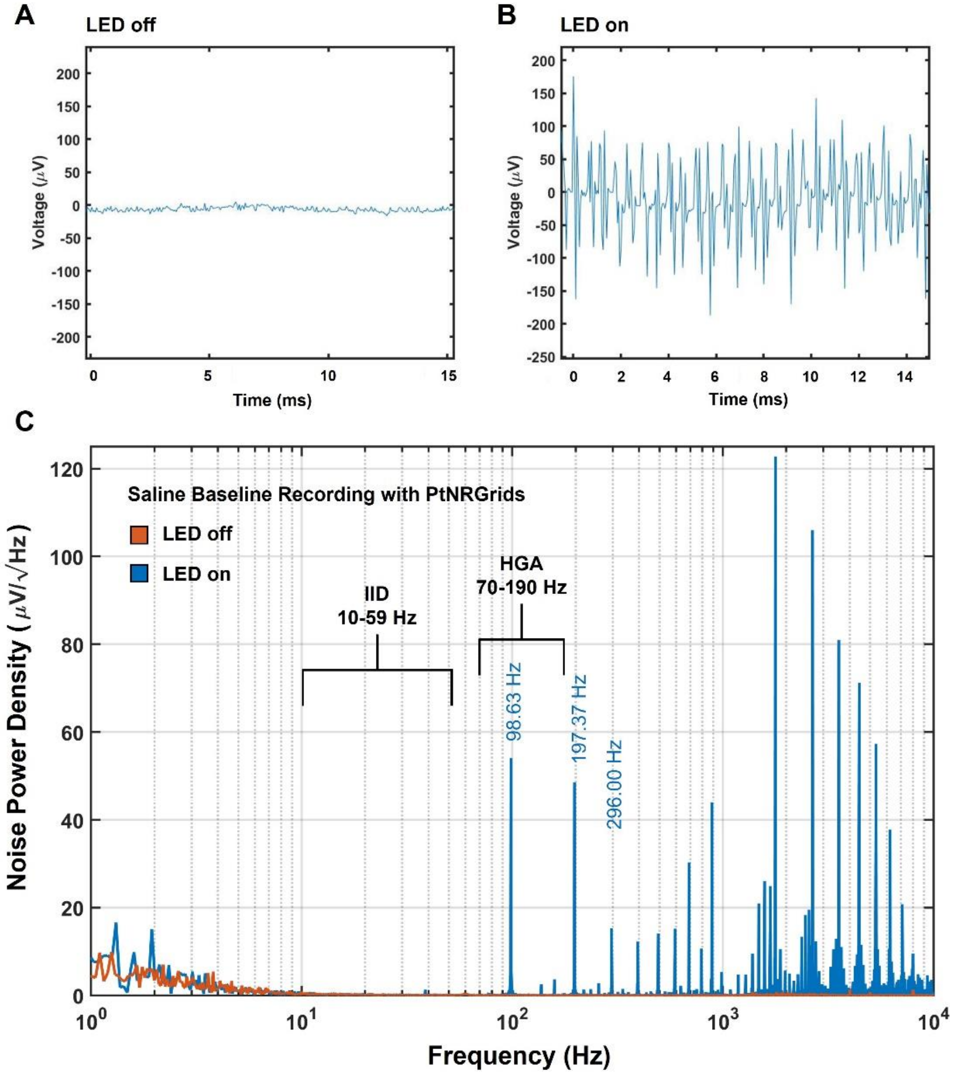


**Figure S10.** Electrical noise induced on µECoG by the µLED array operation. Raw waveform of µECoG grid recording baseline of saline with (A) LED powered off and (B) LED under operation. (C) Noise power density of µECoG signals with and without LED operation. The two main frequency windows, i) 10-59 Hz and ii) 70-190 Hz that we analyzed the data, are marked in the graph.

Table S1. Table summarizing pig models, anesthesia, type of device used, epileptogenic neurotoxin, and summary of recording.

| Date | Breed | Weight (kg) | Anesthesia | Anesthesia Maintenance | Type of Brain EEG-Microdisplay | Epileptogenic neurotoxin | Task |
|---|---|---|---|---|---|---|---|
| Pilot Study | Female Yorkshire | 74 | Midazolam 0.5 mg/kg<br>Ketamine 10 mg/kg<br>Atropine 0.004 mg/kg | Isoflurane and<br>Propofol<br>8mg/kg/hr | 32mm × 32mm, 1024 pixel | - | M1/S1 boundary, HGA mapping |
| Nov 15, 2022 | Female Yorkshire | 62 | Midazolam 0.5 mg/kg<br>Ketamine 10 mg/kg<br>Atropine 0.004 mg/kg | Isoflurane and<br>Propofol<br>8mg/kg/hr | 32mm × 32mm, 1024 pixel | Pen-G | M1/S1 boundary, HGA mapping,<br>Ojemann probe stimulation |
| Nov 16, 2022 | Female Yorkshire | 62 | Midazolam 0.5 mg/kg<br>Ketamine 10 mg/kg<br>Atropine 0.004 mg/kg | Isoflurane and<br>Propofol<br>8mg/kg/hr | 32mm × 32mm, 1024 pixel | Pen-G | M1/S1 boundary, HGA mapping,<br>Ojemann probe stimulation |
| Jan 12, 2023 | Female Yorkshire | 72 | Midazolam 0.5 mg/kg<br>Ketamine 10 mg/kg<br>Atropine 0.004 mg/kg | Isoflurane and<br>Propofol<br>8-10mg/kg/hr<br>Vecuronium<br>5mg bolus<br>2mg/hr IV | 13mm × 32mm, 2048 pixel dual-color | 4-AP | M1/S1 boundary, HGA mapping,<br>IID mapping,<br>Ojemann probe and sEEG stimulation |
| Jan 13, 2023 | Female Yorkshire | 72 | Midazolam 0.5 mg/kg<br>Ketamine 10 mg/kg<br>Atropine 0.004 mg/kg | Isoflurane and<br>Propofol<br>8mg/kg/hr | 13mm × 32mm, 2048 pixel dual-color | BIC | M1/S1 boundary, HGA mapping,<br>IID mapping, Ojemann probe and sEEG stimulation |

Table S2. Table summarizing rat models, anesthesia, type of device used, epileptogenic neurotoxin, and summary of recording.

| Date | Breed | Weight (g) | Anesthesia | Anesthesia Maintenance | Type of Brain EEG-Microdisplay | Epileptogenic neurotoxin | Task |
|---|---|---|---|---|---|---|---|
| Nov 9, 2022 | Male Sprague | 500 | Isoflurane 3-4%, transitioned to | Ketamine/Xylazine (100 mg/kg ketamine / | 5mm × 5mm, 1024 pixel | BIC | IID mapping |

| | | | | | | | |
|---|---|---|---|---|---|---|---|
| | Dawley Rats | | Ketamine/Xylazine | 10 mg/kg xylazine) re-dosed every 20-30 min | | | |
| Nov 7, 2022 | Male Sprague Dawley Rats | 500 | Isoflurane 3-4%, transitioned to Ketamine/Xylazine | Ketamine/Xylazine (100 mg/kg ketamine / 10 mg/kg xylazine) re-dosed every 20-30 min | 5mm × 5mm, 1024 pixel | - | Whisker barrel HGA mapping |
| Oct 31, 2022 | Male Sprague Dawley Rats | 500 | Isoflurane 3-4%, transitioned to Ketamine/Xylazine | Ketamine/Xylazine (100 mg/kg ketamine / 10 mg/kg xylazine) re-dosed every 20-30 min | 5mm × 5mm, 1024 pixel | BIC | Whisker barrel HGA mapping, IID mapping, Temperature rise monitor |
| June 9, 2022 | Male Sprague Dawley Rats | 500 | Isoflurane 3-4%, transitioned to Ketamine/Xylazine | Ketamine/Xylazine (100 mg/kg ketamine / 10 mg/kg xylazine) re-dosed every 20-30 min | 5mm × 5mm, 1024 pixel | - | Whisker barrel HGA mapping |
| June 3, 2022 (2nd rat) | Male Sprague Dawley Rats | 500 | Isoflurane 3-4%, transitioned to Ketamine/Xylazine | Ketamine/Xylazine (100 mg/kg ketamine / 10 mg/kg xylazine) re-dosed every 20-30 min | 5mm × 5mm, 1024 pixel | 4-AP | IID mapping |
| June 3, 2022 (1st rat) | Male Sprague Dawley Rats | 500 | Isoflurane 3-4%, transitioned to Ketamine/Xylazine | Ketamine/Xylazine (100 mg/kg ketamine / 10 mg/kg xylazine) re-dosed every 20-30 min | 5mm × 5mm, 1024 pixel | 4-AP | Whisker barrel HGA mapping, IID mapping |

**Movie S1.** Brain-iEEG microdisplay displaying the extent of the electrical field during cortical stimulation of the pig using Ojemann probe.

**Movie S2.** Brain-iEEG microdisplay displaying the extent of the electrical field during cortical stimulation of the pig using sEEG probe.

**Movie S3.** Brain-iEEG microdisplay showing interictal discharge propagation in red color on pig brain together with the M1/S1 boundary displayed in green color.

**Movie S4.** High-resolution Brain-iEEG microdisplay showing interictal discharge propagation on the rat brain.
